# DEFICIENCY OF THE NUTRIENT SENSOR CPT1c IN SF1 NEURONS DISRUPTS THE ENDOCANNABINOID SYSTEM RESULTING IN COMPROMISED SATIETY AND FUEL SELECTION UPON FAT INTAKE

**DOI:** 10.1101/2024.01.23.576690

**Authors:** A Fosch, DS Pizarro, S Zagmutt, AC Reguera, G Batallé, M Rodríguez-García, J García-Chica, O Freire-Agulleiro, C Miralpeix, P Zizzari, D Serra, L Herrero, M López, D Cota, R Rodríguez-Rodríguez, N Casals

## Abstract

The SF1 neurons of the ventromedial hypothalamus (VMH) are pivotal in governing body weight and adiposity, particularly in response to a high-fat diet (HFD). Previous studies have shown that the activation of SF1 neurons induces satiety, increases energy expenditure, and promotes the preferential use of fats as energy substrate. Furthermore, SF1 neurons are necessary for recovering from insulin-induced hypoglycemia. Here we demonstrate the essential role of the nutritional sensor CPT1c in the activation of SF1 neurons by dietary fats. Mice deficient in CPT1C in SF1 neurons (SF1-*CPT1c*-KO) are unable to adjust their caloric intake during the initial exposure to a HFD. This is associated with an impaired metabolic transition in the liver, muscle, and adipose tissue, despite a normal response to a glucose or insulin challenge. During chronic HFD exposure, SF1-*CPT1c*-KO mice are more prone to obesity and glucose intolerance than controls. CPT1c deficiency in SF1 neurons also leads to alterations in hypothalamic endocannabinoid levels and their metabolism. Our findings posit CPT1C in SF1 neurons as a sensor for dietary fats, regulating satiety responses and nutrient partitioning likely through the modulation of the endocannabinoid system.

## INTRODUCTION

The central regulation of body weight is a complex process that involves intricate neural circuits within the hypothalamic nuclei and extrahypothalamic areas, orchestrating appetite and energy homeostasis. The ventromedial hypothalamus (VMH), recognized not only as a satiety center but also as a crucial regulator of energy balance, influences peripheral glucose homeostasis and lipid metabolism (Fosch et al., 2021). Specifically, VMH electric stimulation triggers hypophagia and energy expenditure (EE), and a decrease in respiratory exchange ratio (RER), indicating a preference for fats over glucose as the primary energy substrate to supply the cellular oxidation rate (Ruffin and Nicolaidis, 1999).

The majority of VMH neurons, particularly those in the dorsomedial and central regions of the VMH, express the nuclear receptor steroidogenic factor 1 (SF1), a transcription factor specific to this nucleus and essential for its development and function (Ikeda et al., 1995; Kim et al., 2011, 2009). Optogenetic and chemogenetic studies have revealed that SF1-positive cells are responsible for most of the intrinsic actions of the VMH, including satiety signaling, promotion of energy expenditure (EE), facilitation of fat mobilization (Viskaitis et al., 2017; Zhang et al., 2020) and recovery from insulin-induced hypoglycemia (Meek et al., 2016). Interestingly, transgenic mice with altered expression of various molecular targets in SF1 neurons often display altered metabolic phenotypes, particularly under exposure to high-fat diet (HFD), but not when fed regular chow, emphasizing a role for SF1 neurons in metabolic adaptation to increased fat consumption (Fosch et al., 2021). Moreover, recent data suggest that unsaturated fats, but not saturated ones, activate SF1 neurons driving satiety and EE (Fosch et al., 2023). However, the molecular mechanisms and neural circuitry that underlie the sensing of fats by SF1 neurons remains poorly understood. Notably, the endocannabinoid (eCB) receptor CB1 in SF1 neurons decreases the excitability of those neurons and modulates feeding and fuel selection by peripheral tissues based on the type of diet (Cardinal et al., 2014; Kim et al., 2008).

The neuron-specific carnitine palmitoyltransferase 1c (CPT1c), emerges as a key player in appetite regulation and nutrient partitioning in the hypothalamus, including the VMH (Dai et al., 2007; N Price et al., 2002). Global CPT1c knock-out (KO) mice exhibit impaired feeding responses to leptin and ghrelin (Gao et al., 2011; Ramírez et al., 2013), and disrupted diet-induced brown fat thermogenesis, making them more susceptible to become obese when fed a HFD (Rodríguez-Rodríguez et al., 2019). Moreover, CPT1c is involved in food preference (Okamoto et al., 2018), fuel selection under fasting conditions (Pozo et al., 2017), and the sensing of fats by the hypothalamus (Fosch et al., 2023). However, there is limited data regarding the role of CPT1c in specific nuclei of the hypothalamus. At the molecular level, CPT1c, unique in lacking enzymatic activity within the CPT1 family (Wolfgang et al., 2006), functions as a nutrient sensor by binding malonyl-CoA within neurons (Casals et al., 2016; Casas et al., 2020; Fadó et al., 2021; Miralpeix et al., 2021a; Palomo-Guerrero et al., 2019; Nigel Price et al., 2002; Rodríguez-Rodríguez et al., 2023). Malonyl-CoA is a lipid metabolite that dynamically fluctuates in the hypothalamus based on the nutritional state (Tokutake et al., 2012, 2010) and regulates feeding behavior and energy homeostasis (Cha et al., 2006; Gao et al., 2007; He et al., 2006; Hu et al., 2003; Lane et al., 2008; López et al., 2006; Wolfgang et al., 2007). Our recent studies have shown that the regulation of glutamate-type AMPA receptors trafficking and synaptic transmission by CPT1c depend on malonyl-CoA (Casas et al., 2020; Fadó et al., 2015; Gratacòs-Batlle et al., 2018). In addition, CPT1c interacts with the eCB hydrolase α/β-hydrolase domain-containing 6 (ABHD6), modulating its enzymatic activity depending on nutrients and malonyl-CoA availability (Miralpeix et al., 2021a). Building upon this evidence, here we have hypothesized a role for CPT1C within SF1 neurons of the VMH in the response to dietary fats. Our findings demonstrate that the specific deletion of CPT1c in SF1 neurons diminishes the responsiveness of SF1 neurons to fats and increased eCBs levels, thereby leading to impaired satiety signaling, disrupted nutrient partitioning, and ultimately favoring obesity.

## METHODS

### Animals

Mice were housed on 12h/12h light/dark cycle (light on at 8 am, light off 8 pm) in a temperature- and humidity-controlled room. Animals were allowed free access to water and standard laboratory chow diet, otherwise indicated. All animal procedures were performed in agreement with European guidelines (2010/63/EU) and approved by the Ethical Committee of the University of Barcelona (n. 10906 and n. 10210 from the Generalitat de Catalunya) and of the University of Bordeaux (n. 30252).

### Generation of SF1-CPT1c-KO mice

Mice deficient of *CPT1c* in SF1 neurons were generated through two consecutive steps using Flp-FRT and Cre-loxP technologies at CBATEG from Universitat Autònoma de Barcelona. Sperm of the strain C57BL/6N-CPT1c^tm1a(EUCOMM)Wtsi^, containing exons 4 to 6 of *Cpt1c* flanked by two LoxP sequences, a lacZ cassette and a neo gene, was obtained from KOMP Repository (University of California, Davis). This sperm was used for an *in vitro* fertilization to generate heterozygous mice called KO-first. LacZ and Neo cassettes were deleted by breeding KO-first with mice containing FlpO in heterozygosis (FlpO mice) from the Mouse Mutant Core Facility (MMCF) of IRB Barcelona, to get a CPT1c^loxP/-^ mice. These animals were bred with SF1-Cre mice (Tg(Nr5a1-cre)Lowl/J, stock number 012462 from The Jackson Laboratory, Bar Harbor, ME) to generate SF1-Cre;*CPT1c*^loxP/-^, which finally were bred with a homozygous *CPT1c*^loxP/loxP^ mice to get SF1-Cre;*CPT1c*^loxP/loxP^, also called SF1-*CPT1c*-KO. *CPT1c*^loxP/loxP^ mice resulting from this last breeding were control littermates called SF1-*CPT1c*-WT mice.

### Genotyping

DNA from VMH or ear punches were used for genotyping. VMH samples were collected using a brain matrix and stored at -80 °C until used. DNA was obtained using Proteinase K (3115887001, Roche, Basel, Switzerland) digestion followed by phenol-chloroform extraction and ethanol precipitation. DNA from ear punches was extracted with 50 mM NaOH and buffered in TRIS 1M pH 8. The oligonucleotide enhancers used for DNA amplification are described in Supplemental Fig. 1.

### Administration of adeno-associated virus (AAVs) within the VMH

For the stereotaxic surgery, mice were anesthetized with 75 mg/kg of ketamine (Ketamidor® 100 mg/mL Ricther Pharma AG, Bagres Austria) and 10 mg/kg of xylazine (Rompun® 20 mg/mL Bayer, Leverkusen, Germany). Stereotaxic surgery to target VMH was carried out with the following coordinates: 1.5 mm posterior from Bregma, ± 0.5 mm lateral to midline and 5.8 mm deep. Purified AAVs expressing mCherry under Cre activity (AAV-CRE-mCherry) (Zagmutt et al., 2023) were injected bilaterally over 10 minutes through a 33-gauge injector connected to a 5 µL Hamilton® Syringe (65460-02, Hamilton Company, Reno, USA) and an infusion pump. 0,4 nL with a viral titer of 1,23x10^12^ pfu/mL were infused in each injection site. After surgery, an analgesic solution (Meloxidyl® 5 mg/mL, Ceva Santé Animale, Libourne, France) was subcutaneously administered to a final dose of 1 mg/kg of body weight. During the first two days of recovery, the analgesic was administered through the drinking water with a final solution of 5 mg/L. Mice underwent 3 weeks of recovery before their use for the study.

### In situ hybridization

RNA Scope® technology (322340 and 320851, ACD Bio, Newark, USA) was used for fluorescent *in situ* hybridization (FISH) detection of *Sf1* and *Cpt1c* mRNA following manufacturer recommendations in fixed brain sections of 15 µm thickness. Then, sections were mounted using antifade Fluoromount-G® (0100-01, Southern Biotech, Birmingham, USA) on coverslips (DIO2460, Deltalab, Barcelona, Spain) and analyzed by LEICA DMi8 confocal microscope.

### Administration of high fat diets

Mice were fed a HFD (60% kcal from fat, ref. D12492) or a standard diet (SD, 10% kcal from fat, ref. D12450J, Research Diets, New Brunswick, USA) for 5 days or 8 weeks. During this period, body weight and the amount of each diet eaten was weakly measured. At the end of the study, mice were sacrificed by cervical dislocation and hypothalamus, epidydimal white adipose tissues (eWAT), subcutaneous WAT (sWAT), brown adipose tissue (BAT), liver and soleus muscle were collected and stored at -80 °C until further processing. For the analysis of neuronal activation by c-Fos, mice were fasted for 3 hdurign the light phase followed by a refeeding period of 2 hours with HFD or SD. Then, mice were perfused for brain cryopreservation.

### Indirect calorimetry, in cage-locomotor activity and gas exchange analysis

Indirect calorimetry, in-cage locomotor activity and gas exchange analysis were carried out in light, temperature and humidity controlled calorimetric chambers (TSE Systems GmbH, Moos, Germany) as described previously (Castellanos-Jankiewicz et al., 2021). The light cycle was 12 h/12 h light/dark phase (lights on 3am, lights off 3pm) at 22 °C. Mice were acclimated for 5 days before recording. O2 consumption and CO2 production were measured to calculate RER (or respiratory quotient, RQ) and EE. Locomotor activity was determined using an infrared light beam system. All measurements were taken every 20 minutes.

### Glucose and insulin tolerance tests

All glucose measurements were performed using the glucometer Aviva from Accu-Chek® (Roche, Basel, Switzerland) and test strips (06453970, Roche, Basel, Switzerland) on the second drop after a little cut on the tail. Glucose tolerance test (GTT) and insulin tolerance test (ITT) were conducted in 6 h fasted mice, starting at the beginning of the light phase. 20 % glucose (Glucose 20 % B. Braun Medical, Melsungen, Germany) was injected intraperitoneally with a final dose of 2 g/kg body weight. 0.1 IU/mL of human insulin (Humulin 100 IU/mL, Lilly Medical, Indianapolis, USA) was injected with a final dose of 0.5 IU/kg body weight. Data were represented as glycaemia evolution over time and area under the curve. Glucose disappearance rate (K_ITT_) was calculated using the first 30 min.

### Leptin sensitivity test

Mice were weighed and fasted 1 hour before intraperitoneal injection. PBS (vehicle) or 0.5 mg/mL of leptin (498-OB-05M, Bio-Techne Sales Corp, Minneapolis, USA) with a final dose of 2.5 mg leptin/kg were injected during the lightt cycle. Food intake was measured at 1 h, 4 h and 24 h. All mice received vehicle injection the day before leptin injection.

### Intracerebroventricular (ICV) cannulation surgery

Cannulas were stereotaxically implanted into the lateral cerebral ventricle, being the coordinates 0.58 mm posterior to Bregma, 1 mm lateral to the midsagittal suture and 2.2 mm of depth. Intraperitoneal ketamine (75 mg/kg body weight) plus xylazine (10 mg/kg body weight) was used as anesthesia. Mice were individually caged and allowed to recover for at least 5 days before the experiment. Cannula placement was verified by a positive dipsogenic response to angiotensin II (1 nmol in 1 mL; Sigma-Aldrich, Saint Louise, USA).

### ICV administration of oleic acid

Oleic acid (O3880; Sigma-Aldrich, St Louis, USA) or vehicle were injected into ICV-cannulated mice. Oleic acid was complexed with fresh 40 % HPB (2-Hydroxypropyl)-β-cyclodextrin (H107 Sigma-Aldrich, St Louis, USA). 9 nmols of oleic acid/dose and 2 µl/dose were injected. Animals did not have access to food during the whole procedure. After 2 h, mice were perfused for brain collection, as indicated below.

### RNA preparation and quantitative RT-PCR

Total RNA was extracted from tissues using Trizol Reagent (Fisher Scientific, Madrid, Spain). Quantitative RT-PCR analysis was performed as previously described (Fosch et al., 2023). SsoAdvanced^TM^ SYBR®Green Supermix (172-5261, BioRad, Hercules, USA) was used for double-stranded DNA intercalating agents and SsoFast^TM^ Probes Supermix (172-5231, BioRad, Hercules, USA) for TaqMan assay. SYBR Green assay primers are listed in Supplemental Fig 2. Taqman Gene Expression assay primers for *Ucp1* were used (for: 5’-CACACCTCCAGTCATTAAGCC, and rev: 5’-CAAATCAGCTTTGCCTCACTC). All primers were from IDT DNA Technologies, Leuven, Belgium. Relative mRNA levels were measured using the CFX96 Real-time System, C1000 Thermal Cycler (BioRad, CA, USA). Relative gene expression was estimated using the comparative Ct (2^-ΔΔCt^) method in comparation to *Gapdh* levels.

### Western blotting

Western blot was performed as previously described (Fosch et al., 2023). Briefly, soleus muscle was homogenized in RIPA buffer (Sigma-Aldrich, Madrid, Spain) containing protease and phosphatase inhibitor cocktails. Protein extracts were separated on SDS-PAGE, transferred into Immobilion-PVDF membranes (Merck Millipore, Madrid, Spain) and probed with antibodies against: ACC, pACC (Ser79) (Cell Signaling; Danvers, MA, USA); and GAPDH (Abcam, Cambridge, UK). Anti-mouse or anti-rabbit horseradish peroxidase-conjugated (Jackson, West Grove, USA) were used as secondary antibodes. LuminataForte Western HRP substrate (Merck Millipore, MA, USA) was used as developing agent. Images were collected by the ChemiDoc MP and Image Lab Software (Bio-Rad Laboratories, Hercules, USA) and quantified by densitometry using ImageJ-1.33 software (NIH, Bethesda, MD, USA). In all the figures showing images of gels, all the bands for each picture come from the same gel, although they may be spliced for clarification.

### Brain immunofluorescence

Mice were anesthetized under ketamine/xylazine and intracardially perfused with PBS and then 10 % NBF. Brains were collected and postfixed 24 h in 10 % NBF at 4 °C, transferred to 30 % sucrose at 4 °C for 2-3 days, frozen in isopentane, and sliced in 30 µm thick slices in the coronal plane throughout the entire rostral-caudal extent of the brain using a cryostat. Slices were preserved at -20 °C in antifreeze solution until use.

Sections containing the hypothalamus were processed as previously described (Fosch et al., 2023). Briefly, sections were blocked in 2 % goat or donkey anti-serum in KPBS and 3 % BSA plus 0.1 % Triton X-100 and incubated with primary antibody (rabbit anti-c-Fos, 1:200, Cell Signaling, Danvers, MA, USA; sheep anti-αMSH, 1:1000, Sigma-Aldrich, Merck KGaA, Darmstadt, Germany) for 1 h at room temperature or overnight at 4 °C, respectively. Goat anti-rabbit Alexa Fluor 647 antibody (1:1000; Invitrogen, Waltham, USA) or donkey anti-sheep Alexa Fluor 488 (1:1000; Invitrogen, Waltham, USA) were used as secondary antibodies. Slices were counterstained with Hoechst for nuclear staining (1 mg/mL, Sigma-Aldrich, Saint Louis, USA). Then, slices were mounted using antifade Fluoromount-G® (0100-01, Southern Biotech, Birmingham, USA) on coverslips (DIO2460, Deltalab, Barcelona, Spain). Images were taken using a Leica DMi8 confocal microscope equipped with a 20x and 40x objective. Fluorescence integrated density after image masking was calculated using ImageJ 1.33 software.

### Adipose tissue histopathology and adipocyte area measurement

Fixed eWAT were dehydrated and embedded in paraffin. The resulting blocks were cut into 5–10 µm sections and stained with hematoxylin and eosin (H&E) to assess histology. To quantify the adipocyte area, three representative images from each adipose tissue section were taken with a 20x objective using a high-sensitivity camera (Leica MC 190 HD Camera) and microscope. Images were analyzed with the Adiposoft software as described in Zagmutt et al., 2023. Images were analyzed and adipocytes were highlighted if they met the following criteria: (1) the boundaries for sizing of the cell were 40-40,000; (2) the adipocyte had a shape factor of 0.35–1 (a shape factor of 0 indicated a straight line, while a shape factor of 1 indicated a perfect circle); and (3) the adipocyte did not border the image frame. The results are represented as the average of total area counted.

### Oil Red-O liver tinction

Livers were collected and fixated 24 h in 10% NBF at 4°C, transferred to 30% sucrose at 4°C for 2-3 days, frozen in isopentane, and sliced in 30 µm thick slices using a cryostat. Slices were mounted, fixated with 10% NBF and dehydrated with 90% isopropanol, and then stained with Oil Red-O. Images were obtained with 10x objective using a high-sensitivity camera (Leica MC 190 HD Camera) and quantified using ImageJ-1.33 software.

### Extraction and analysis of eCBs

Hypothalamic eCBs were extracted and analyzed as described in Miralpeix et al., 2019. Hypothalamus (6–8 mg wet tissue) was homogenized in in a Dounce tissue grinder with 200 μL of ice-cooled deionized water containing a final concentration of 0.362 μM N-oleylethanolamine-d2 (OEA-d2) (Cayman Chemicals, Ann Arbor, MI) as the internal standard for 2-arachydonoyl (2-AG) and anandamide (AEA) calibration, 100 μM PMSF, and 0.01 % butylated hydroxytoluene (BHT) (Sigma-Aldrich, Madrid, Spain), followed by a brief sonication. After that, lipids were extracted with ethyl acetate/n-hexane (9:1, v/v) and evaporated. eCBs levels were analyzed by LC/MS/MS. 2-AG, AEA, and OEA-d2 were used for the calibration curve in an Acquity ultra-high-performance liquid chromatography (UPLC) (Waters, Singapore) system connected to a Xevo-TQS triple-quadropole Detector (Waters, Ireland) and controlled with Waters/Micromass MassLynx software. Chromatographic separation was performed on an Acquity UPLC BEH C18 column with an isocratic mobile phase of formic acid 0.1 % in water-acetonitrile (30:70, v/v). Detection was performed in the positive ion mode. The capillary voltage was set to 3.1 kV, the source temperature was 150 °C, and the desolvation temperature was 500 °C.

### Tissue triglycerides and serum leptin determination

40 mg of tissue (liver or WAT) were homogenized in cold acetone using ceramic beads (32201-1L-M, Sigma-Aldrich, Saint Louis, USA) and Fast-Prep®-24 at 6.5 M/sec for 3 cycles of 30 seconds. Then, samples were incubated in rotatory homogenization o/n at RT. Upper layer was used for TG measurement using Serum Triglyceride Determination kit (TR0100, Sigma Aldrich, Saint Louis, USA). Serum leptin was measured with Mouse Leptin ELISA KIT #90030, Crystal Chem, Grove Village, IL, USA.

### Data processing and statistical analysis

Statistical analyses were performed with Prism 9.0 (Graphpad Software, San Diego, USA). Data were expressed as mean ± SEM. Two-way ANOVA test followed by post hoc two-tailed Bonferroni test was applied or a t-student test if only two groups were compared. A p-value less than 0.05 (p<0.05) was considered statistically significant. The mice number and the statistical test used are specified in each figure legend.

## RESULTS

### SF1-*CPT1c*-KO mice show increased food intake, respiratory quotient and adiposity under short-term high-fat feeding

We generated a mouse model with specific deletion of CPT1c in SF1 neurons using the FRT-Flp and Cre-lox recombination technologies as described in the methodology section. We confirmed that SF1-*CPT1c*-KO mice were properly generated by 1) the presence of Cre recombinase activity at the VMH (Fig. 1A), 2) the amplification of a 531 bp band containing the deletion of exons 4 to 6 of *CPT1c* only when DNA was obtained from VMH but not from hippocampus or cortex brain samples (Fig. 1B), and 3) the negligible expression of CPT1c in SF1 neurons measured by RNAScope® FISH (Fig. 1C)

**Fig. 1.**
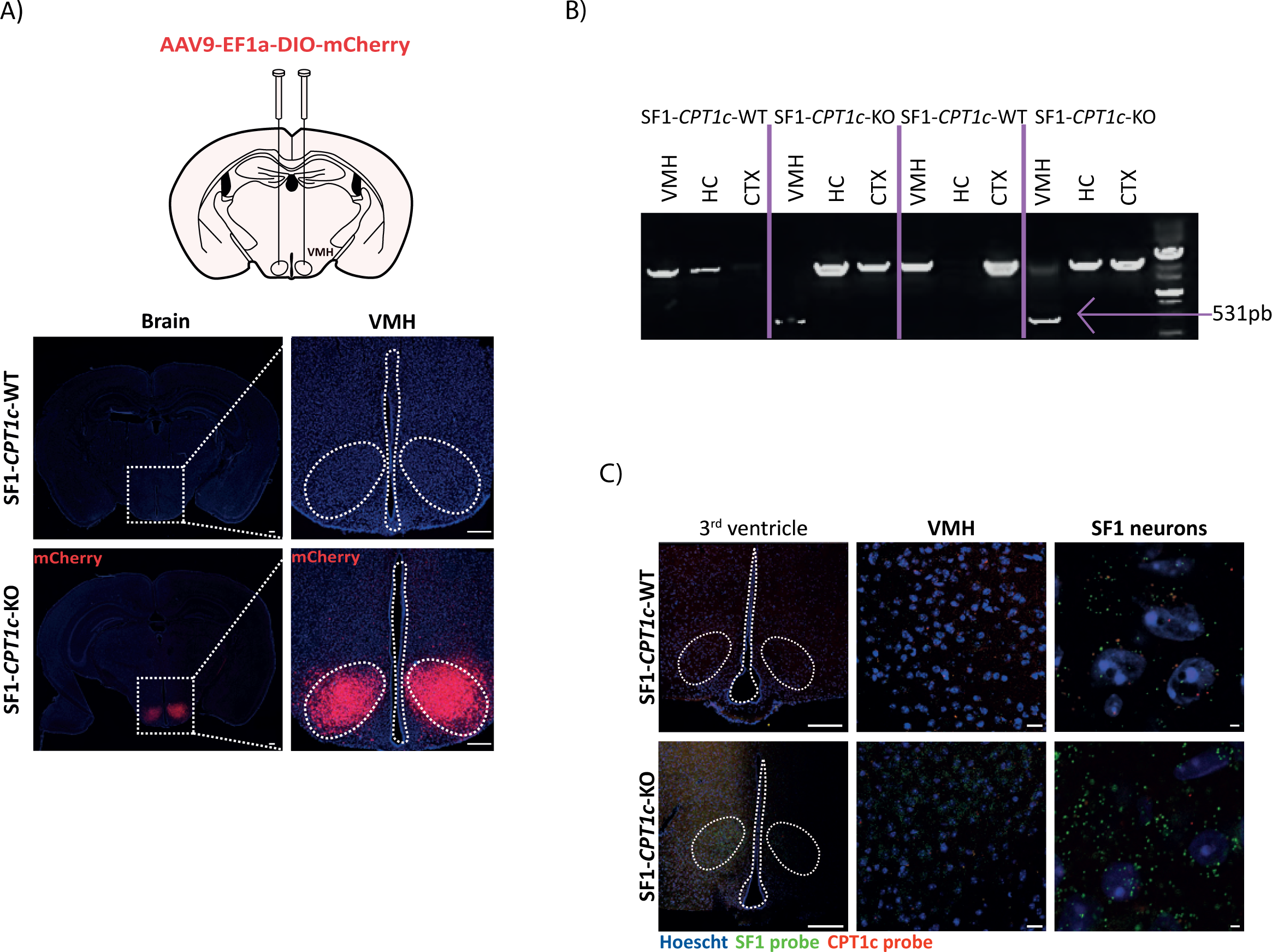
Validation of SF1-*CPT1c*-KO mice. A) Validation of CRE activity of SF1 cells. Upper, schematic representation of stereotaxic injection of AAV-DIO-mCherry (AAVs that express mCherry under the activity of CRE); below, VMH area in brain sections from SF1-*CPT1c*-WT and SF1-*CPT1c*-KO mice, 3 weeks after AAVs injection. Scale bar: 250 µm. Only SF1-*CPT1c-*WT show CRE activity in the VMH. B) PCR amplification to demonstrate the excision of the exons 4 to 6 of the gene *CPT1c* in SF1-*CPT1c*-KO mice. Picture of agarose gel after PCR product was run. Samples derived from three different parts of the brain were used. CTX: cortex; HC: hippocampus; VMH: ventromedial hypothalamus. A lower-sized band corresponding to the excision of exons 4-6 is only present in the VMH of SF1-*CPT1c*-KO mice. C) *Sf1* and *CPT1c* expression in VMH using RNAScope® FISH technique. Probes against SF1 mRNA (green) and CPT1c mRNA (red) were designed and used in brain sections obtained from SF1-*CPT1c*-WT and -KO mice. From left to right: 3^rd^ ventricle, VMH and zoom of VMH area (SF1 neurons). Scale bar 3^rd^ ventricle: 250 µm; VMH: 25 µm; SF1 neurons: 5 µm.

For the phenotyping, we first monitored the evolution of feeding and body weight of male and female mice over 8 weeks of SD and found no differences between genotypes (Supplemental Fig. 3-5). Neither adiposity, EE, RER, locomotor activity, brown adipose tissue (BAT) thermogenesis, glucose tolerance, insulin or leptin sensitivity, nor leptin-induced thermogenesis were altered between SF1-*CPT1c*-KO and SF1-*CPT1c*-WT.

Interestingly, switching animals to a HFD (60 % of calories from fat) for just five days was enough to reveal significant differences in crucial parameters between genotypes. On the one hand, male SF1-*CPT1c*-KO mice showed higher food intake, especially during the dark phase (Fig. 2A-B), leading to an increase in body weight (Fig. 2C). As illustrated in Fig. 2D, SF1-*CPT1c*-KO mice do not properly accommodate total caloric intake upon dietary change to fats as control animals do. On the other hand, calorimetric studies showed that the RER of SF1-*CPT1c*-KO was higher than that of control mice, especially during the dark phase (Fig. 2E). This implies that in KO animals subjected to HFD, there is a reduced inclination to utilize fats as the primary fuel source in peripheral tissues. Magnetic Resonance Imaging (MRI) analysis revealed an increase of fat mass (Fig. 2F). The increase in fat mass was not due to reduced energy expenditure, BAT thermogenesis or locomotor activity, nor to increased feed efficiency (Supplemental Fig. 6). Leptin levels remained similar between genotypes (Fig. 2G).

**Fig. 2.**
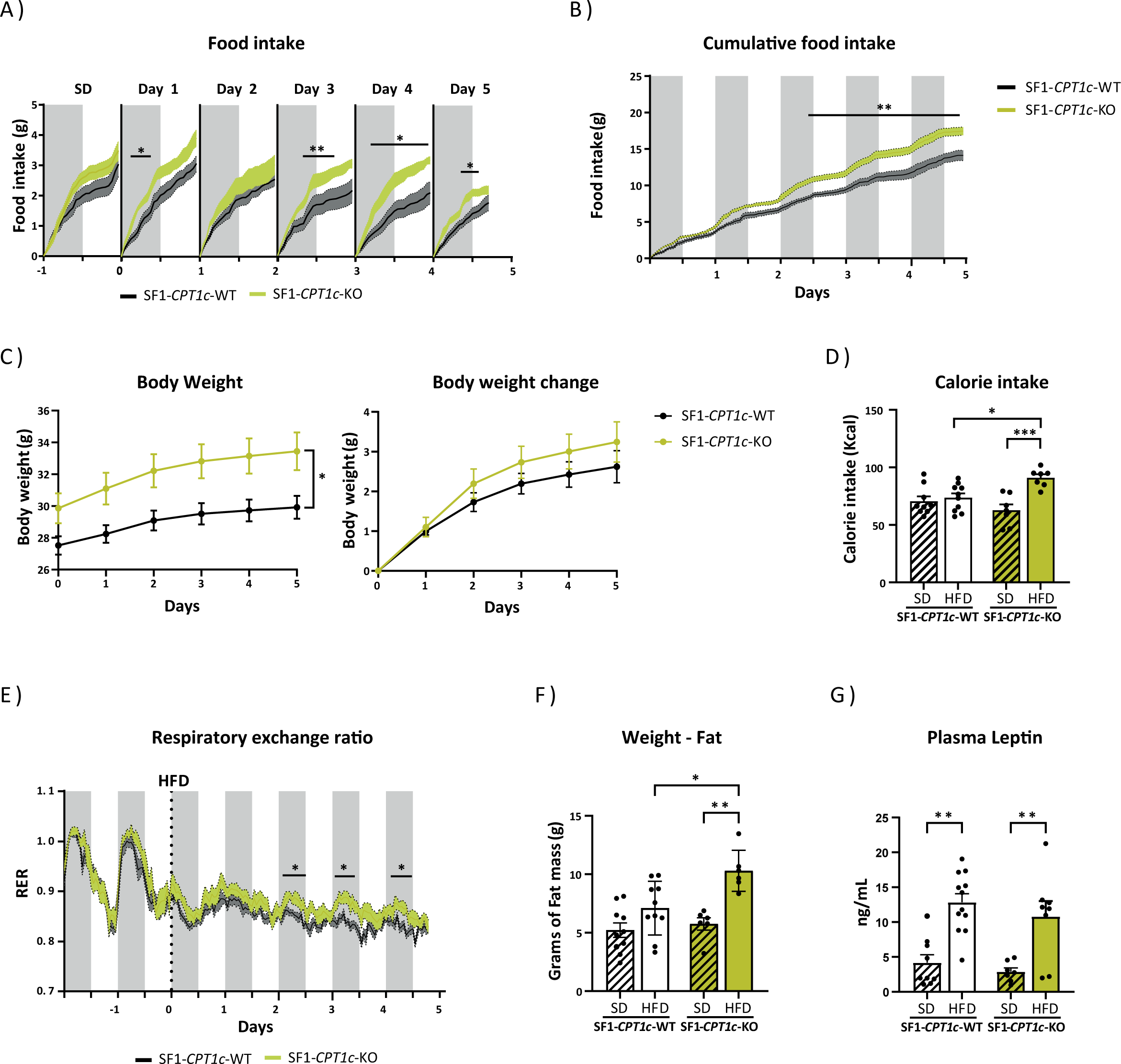
Metabolic phenotype of SF1-*CPT1c*-KO and -WT mice upon 5 days of HFD. A) Cumulative daily food intake was recorded during both the dark and light phases of each day, starting one day prior to diet change and continuing for 5 days while on the high-fat diet (HFD). B) Cumulative food intake. C) Body weight and body weight gain. D) Total calorie intake over the 5 days of HFD. E) Respiratory exchange ratio, starting 2 days prior to diet change and continuing for 5 days while on the HFD. F) Magnetic Resonance Imaging (MRI) analysis of lean and fat mass of mice in SD and 5 days of HFD. G) Plasma leptin levels. Data are represented as mean ± SEM, male mice, 8-12-week-old, n= 7-10/ group. *p<0.05, **p<0.01, ***p<0.001 vs SF1-*CPT1c*-WT mice. Statistical significance was determined by ANOVA test with post-hoc Bonferroni.

### Peripheral metabolism of SF1-*CPT1c*-KO mice under HFD suggest impaired nutrient partitioning

Next, we analyzed lipid metabolic enzymes in peripheral tissues like liver, white adipose tissue (WAT) and muscle in animals fed HFD for 5 days. qPCR studies showed that liver lipolytic enzymes, such as adipose triglyceride lipase (ATGL) and hormone-sensitive lipase (HSL), and the rate-limiting enzyme of fatty acid oxidation, CPT1A, were decreased while the lipogenic stearoyl-CoA desaturase 1 (SCD1) was increased (Fig. 3A). In relation to lipoprotein lipase (LPL) and CD36, two markers associated with lipid uptake and tissue triglyceride (TG) accumulation (Wang and Eckel, 2009) they increased during the diet switch (after 5 days of HFD) in SF1-*CPT1c*-KO mice but not in control littermates (Fig. 3B-C). Conversely, fatty acid synthase (FASN) expression decreased in both genotypes (Fig. 3D). Collective data suggest an enhanced storage of lipids in liver of SF1-*CPT1c*-KO mice, attributed to increased fatty acid uptake combined with reduced rates of lipolysis and fatty acid oxidation. Accordingly, liver TG and Oil Red-stained fat depots were higher in SF1-*CPT1c*-KO mice than in control mice after only 5 days of HFD (Fig. 4A-B).

**Fig. 3.**
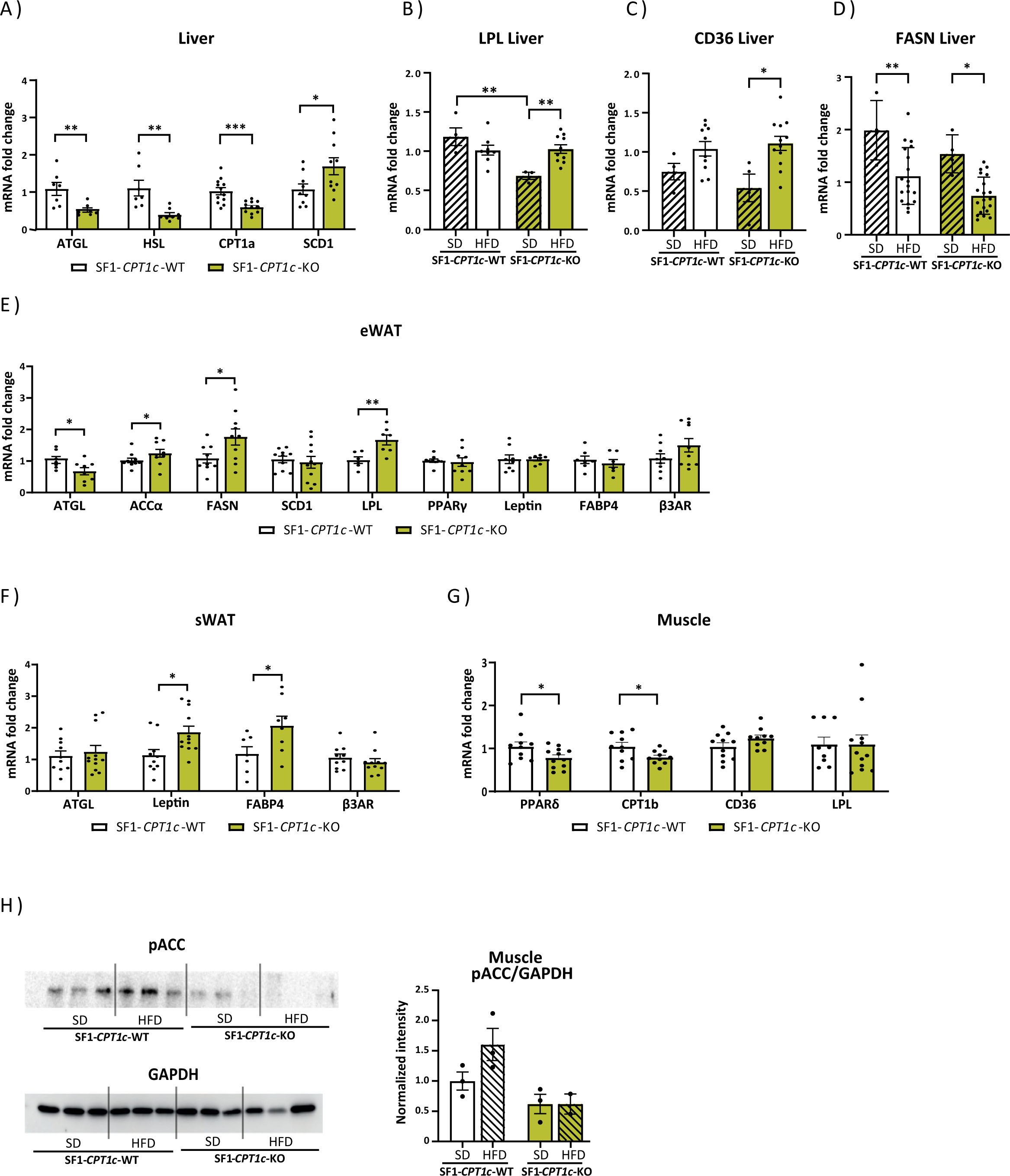
Expression of enzymes, hormones and proteins involved in lipid metabolism in different tissues of SF1-*CPT1c*-KO and -WT mice after 5 days of HFD. A) Expression of adipose triglyceride lipase (ATGL), hormone-sensitive lipase (HSL), CPT1a, stearoyl-CoA desaturase (SCD1) measured by qPCR in liver samples of mice fed a HFD for 5 days. B) Lipoprotein lipase (LPL) expression in liver samples. C) CD36 expression in liver samples. D) Fatty acid synthase (FAS) expression in liver samples. E) Expression of ATGL, acetyl-CoA carboxylase-alpha (ACCa), FAS, SCD1, LPL, peroxisome proliferator-activated receptor gamma (PPARγ), leptin, fatty acid-binding protein 4 (FABP4) and beta-3 adrenergic receptor (b3AR) in epididymal WAT (eWAT). F) ATGL, leptin, FABP4 and b3AR expression in subcutaneous WAT (sWAT). G) PPARδ, CPT1b, CDC36 and LPL expression in gastrocnemius muscle. H) Representative western blot of pACC, ACC and GAPDH from muscle samples and the quantification of pACC normalized by GAPDH. Data are represented as mean ± SEM, male mice, 8-12-week-old, n= 7-12/ group, *p<0.05, **p<0.01, ***p<0.001. Statistical significance was determined by ANOVA test with post-hoc Bonferroni.

**Fig. 4.**
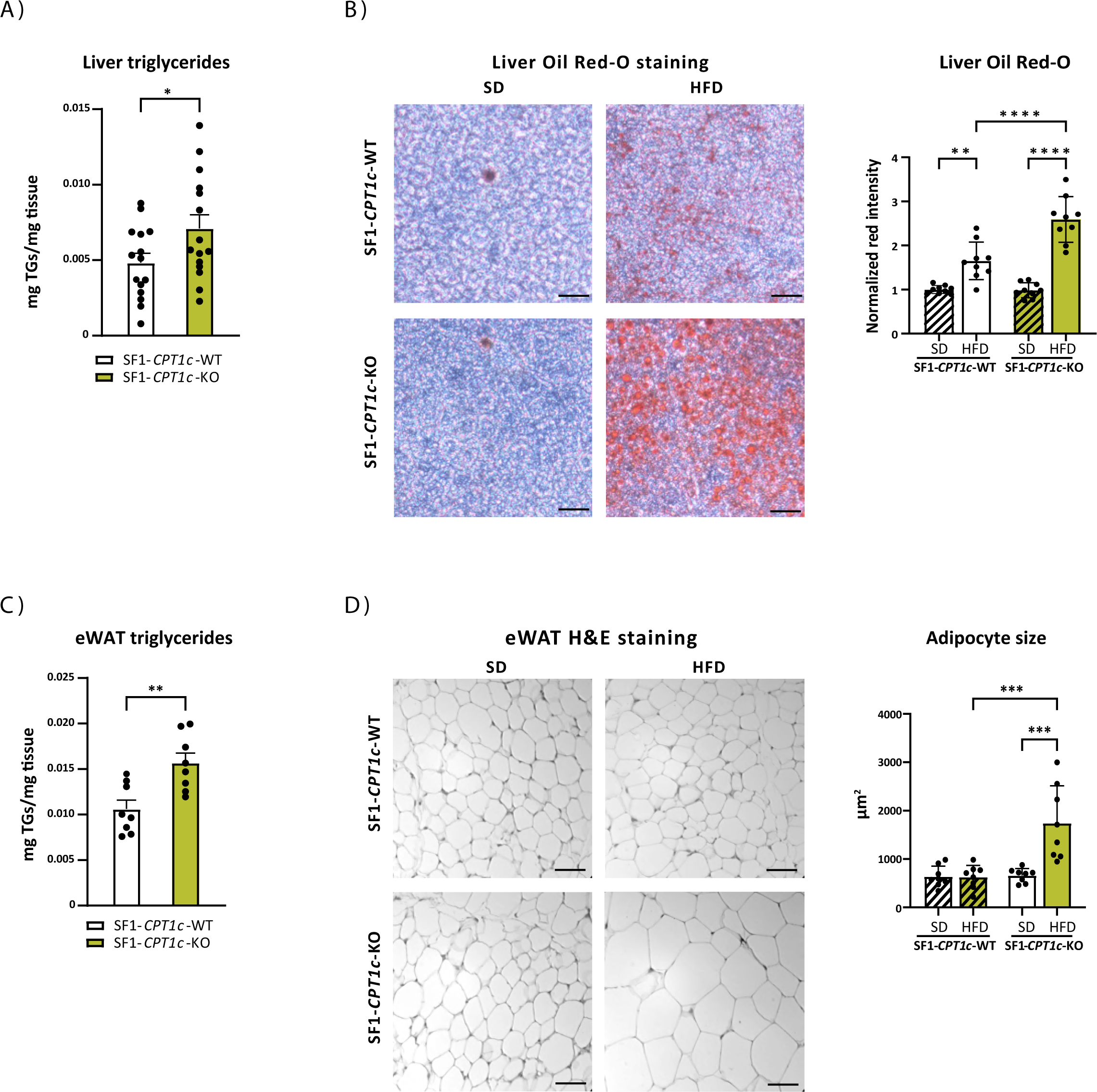
Liver and eWAT adiposity of SF1-*CPT1c*-KO and -WT mice after 5 days of HFD. A) Liver triglycerides after 5 days of HFD B) Oil Red-O staining of liver sections from SF1-*CPT1c*-KO and - WT mice fed a SD or a HFD for 5 days, and quantification. Scale bar: 100µm. C) eWAT triglycerides. D) Hematoxylin and eosin staining of eWAT sections and quantification of adipocyte size. Scale bar: 50µ. Data are represented as mean ± SEM, male mice, 8-12-week-old, n= 8-9/ group, *p<0.05, **p<0.01, ***p<0.001. Statistical significance was determined by ANOVA test with post-hoc Bonferroni.

Similar results were observed in epididymal WAT (eWAT): ATGL was decreased while lipogenic markers such as ACCa and FAS were increased (Fig. 3F), resulting in TG accumulation and larger adipocyte size (Fig. 4C-D). In subcutaneous WAT (sWAT), ATGL levels remained unchanged, but the expression levels of two adipokines secreted during adipogenesis and associated with obesity, namely leptin and FABP4, were higher in KO mice than in WT animals (Fig. 3F). No variations were found in the expression of beta-3 adrenergic receptor (b3AR) in either eWAT or sWAT, in accordance with no changes in EE.

Differences between genotypes were also evident in the soleus muscle. SF1-*CPT1c*-KO mice exhibited lower expression levels of PPARδ and CPT1B (Fig. 3G), along with reduced phosphorylation levels of ACC, in comparison to control animals (Fig. 3H). No changes in CDC36 or LPL were observed. Given that PPARδ serves as a transcription factor that directs cellular energy utilization towards FAO (Liu et al., 2018), CPT1B acts as a primary regulator of FAO in muscle cells, and ACC phosphorylation triggers the activation of CPT1B, these findings suggest that the muscle tissue of SF1-*CPT1c*-KO mice utilizes fats to a lesser extent as an energy source compared to the control mice. Taken together, the findings suggest impaired nutrient partitioning in these mice, as suggested by the higher respiratory quotient.

### The melanocortin signaling is hindered and the sensing of fats blunted in SF1-*CPT1c*-KO mice

To further explore the hyperphagic response observed in SF1-*CPT1c*-KO mice in the first days after switching animals from SD to HFD, the hypothalamic expression of the orexigenic and anorexigenic peptides was analyzed. Results showed that the mRNA expression of the satiating neuropeptide POMC was decreased while the orexigenic AgRP was increased in SF1-*CPT1c*-KO compared to control mice (Fig. 5A). Accordingly, a-MSH staining at the ARC and PVN was decreased in SF1-*CPT1c*-KO mice, specifically upon HFD (Fig. 5B-C), suggesting an impaired satiating response.

**Fig. 5.**
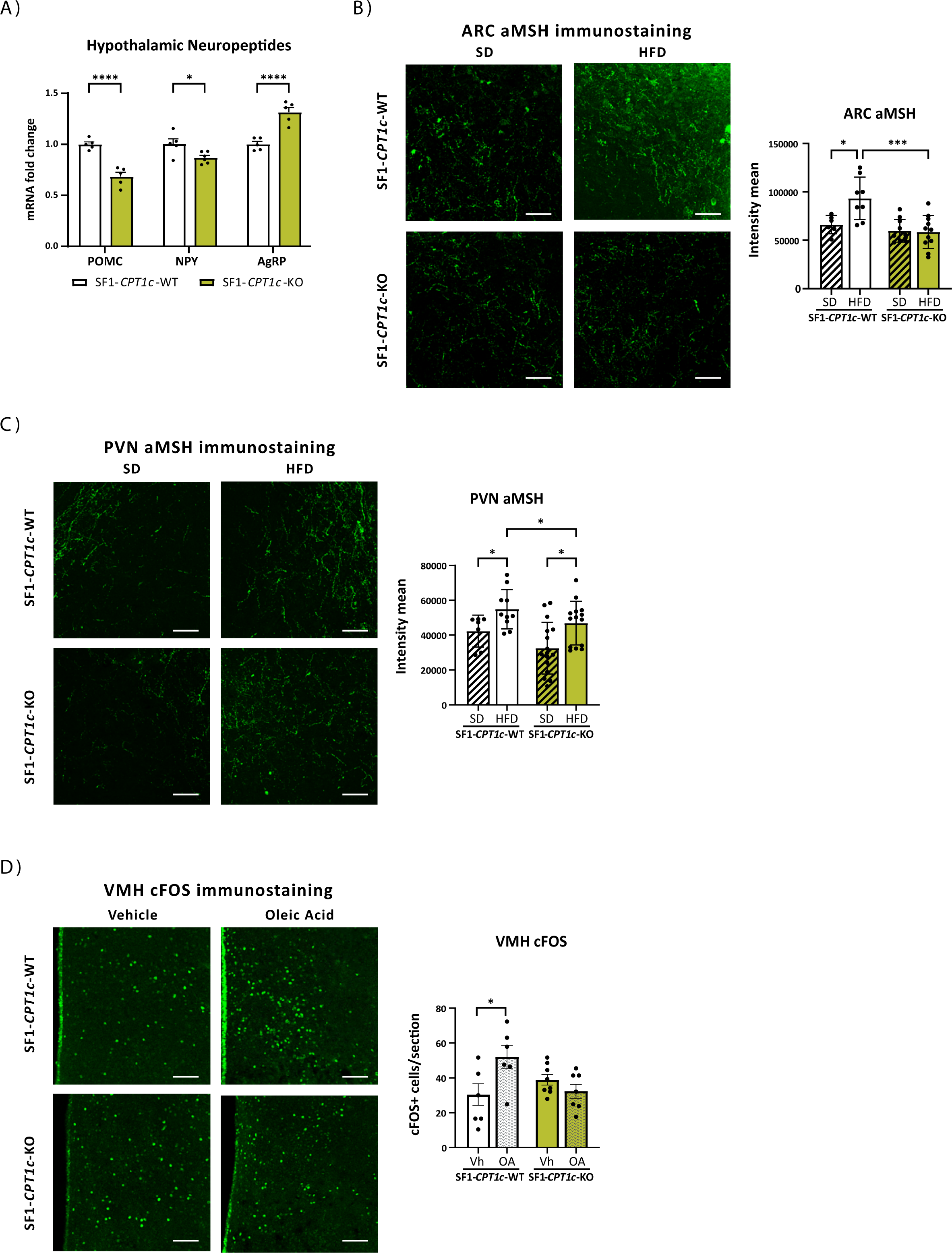
Hypothalamic neuropeptides, alpha-melanocyte stimulating hormone (αMSH), and oleic acid sensitivity. A) Hypothalamic expression of POMC, NPY and AgRP neuropeptides of SF1-*CPT1c*-KO and -WT mice after 5 days of HFD. B) αMSH staining in the PVN. Representative images (left) and quantification of all data (right). C) αMSH staining in the ARC. Representative images (left) and quantification of all data (right). Data from A-C are represented as mean ± SEM, male and female mice, 8–12-week-old, n= 4-5/ group (2-3 slices/mouse). *p<0.05 vs SF1-*CPT1c*-WT. Statistical significance was determined by ANOVA test with post-hoc Bonferroni. D) Effects of oleic acid on neuronal activation assessed by cFOS expression in the VMH. Representative images and quantification of c-Fos positive cells per section. Data are represented as mean ± SEM, male mice, 8-12 weeks of age, n= 3/group (2-3 slices/mouse). Statistical significance was determined by ANOVA test with post-hoc Bonferroni. Scale bar: 100µm.

Since fatty acids are known to signal satiety, and lipid metabolism changes through their action on the hypothalamus (Fosch et al., 2023), we decided to determine whether SF1-*CPT1c*-KO had a defect in FA sensing at the VMH. We injected oleic acid ICV in male mice fed a SD and measured cFOS activation after 2 h. In control mice, oleic acid injection triggered the activation of the VMH (Fig. 5D); however, this was not observed in SF1-*CPT1c*-KO animals. Those findings indicate that CPT1c in SF1 neurons is necessary for proper sensing of fats and activation of VMH nuclei.

### Impaired eCBs levels and metabolism in the hypothalamus of SF1-*CPT1c*-KO mice

Then, we decided to measure eCB levels in the hypothalamus since we have recently demonstrated that they highly increase after short HFD exposure (Miralpeix et al., 2019). Results show that SF1-*CPT1c*-KO mice have much higher levels of both eCBs, anandamide (AEA) and 2-arachidonoylglycerol (2-AG), in the hypothalamus when fed a SD (Fig. 6A). After five days of HFD, control animals increased 2-AG, as previously described (Miralpeix et al., 2019), resulting in similar levels of this eCB in both genotypes. Regarding AEA, although both genotypes exhibit reduced levels after five days of HFD compared to SD, those detected in KO mice consistently remained elevated compared to control mice. Notably, no differences were observed between diets or genotypes in the hippocampus (Supplemental Fig. 7), suggesting that this alteration is specific to the hypothalamus.

**Fig. 6.**
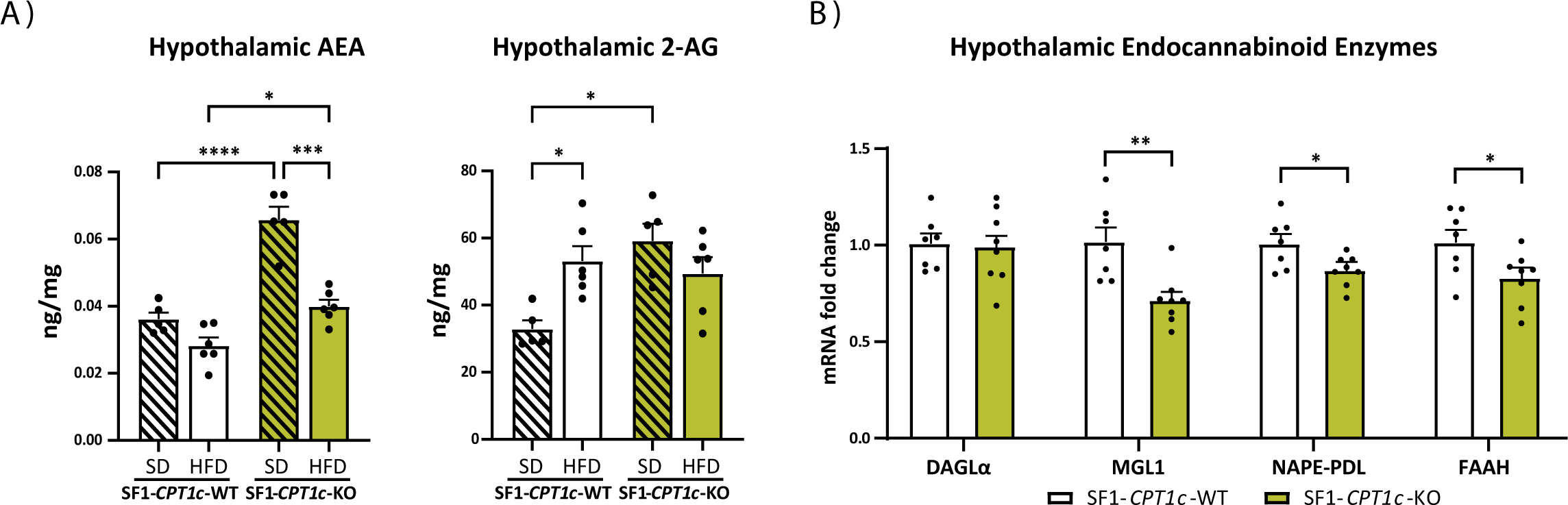
Endocannabinoid system in hypothalamus of SF1-*CPT1c*-KO and -WT mice. A) Hypothalamic levels of anandamide (AEA) and 2-arachidonoylglycerol (2-AG). B) Hypothalamic expression of enzymes involved in the eCB system: diacylglycerol lipase alpha (DAGLα), monoacylglycerol lipase 1 (MGL1), N-acylphosphatidylethanolamine phospholipase D (NAPE-PLD) and fatty acid amide hydrolase (FAAH). Data are represented as mean ± SEM, male and female mice, 8-12-week-old, n= 8-10/ group. *p<0.05 vs SF1-*CPT1c*-WT. Statistical significance was determined by ANOVA test with post-hoc Bonferroni.

Then, we analyzed the expression of eCBs synthesis and metabolizing enzymes in the hypothalamus. Results showed that FAAH and MGL1 expression, the main metabolizing enzymes of AEA and 2-AG, respectively, decreased in SF1-*CPT1c*-KO mice fed a HFD (Fig. 6B), pointing to these metabolic enzymes as responsible for eCBs accumulation. Regarding the synthesizing enzymes, NAPE-PLD exhibited a decrease, suggesting a compensatory response, whereas no changes were observed for DAGLα. In summary, KO animals have elevated baseline levels of eCBs under SD conditions and exhibit an anomalous response in their dynamics when switched to a HFD.

### SF1-*CPT1c*-KO mice develop obesity and exhibit a diabetic phenotype over the long term HFD exposure

Maintaining the mice on a HFD for a longer period (8 weeks) resulted in greater differences between genotypes in adiposity and body weight than the ones observed in 5 days of HFD, being the differences more prominent in male animals compared to female mice (Fig. 7A-G and Supplemental Fig. 8A-D). Notably, in the long term HFD feeding, the amount of food consumed was the same in both genotypes (Fig. 7B and Supplemental Fig. 8C). However, alterations in the expression of some lipid metabolism markers in peripheral tissues were maintained (Fig. 7F) suggesting that the increased adiposity observed in the KO mice is primarily due to impaired nutrient partitioning. Interestingly, after 8 weeks of HFD feeding, SF1-*CPT1c*-KO mice showed higher blood leptin levels (Fig. 8A) and impaired insulin sensitivity and glucose tolerance compared to control animals in both male (Fig. 8B-C) and female mice (Supplemental Fig. 8E-F), suggesting the establishment of insulin resistance. In animals fed the SD, although no differences were observed between genotypes regarding the weight of the eWAT, the size of the adipocytes was slightly larger in KO than in WT mice (Fig. 7G), suggesting that with age, they tend to accumulate more fat even on the SD. In summary, the absence of CPT1c specifically in SF1 neurons leads to heightened levels of hypothalamic endocannabinoids and compromised sensing of fatty acids in the VMH. This disruption, in turn, adversely influences the short-term accommodation of caloric intake, along with impaired nutrient partitioning and lipid allocation. These cumulative effects contribute to a heightened susceptibility to an obesogenic and diabetic phenotype over the long term.

**Fig. 7.**
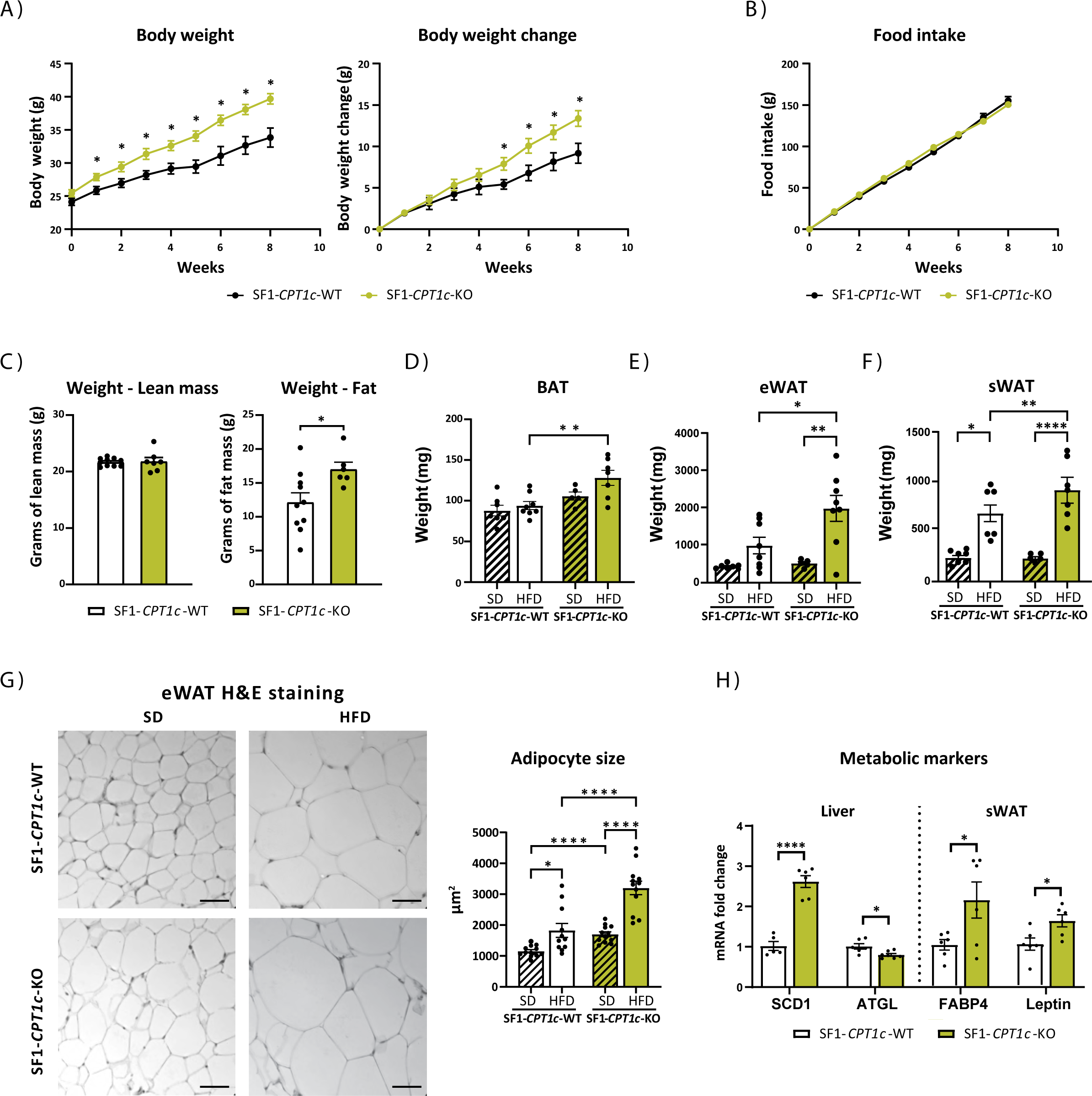
Metabolic phenotype of SF1-*CPT1c*-KO and -WT mice after 8 weeks of HFD. A) Body weight and body weight gain. B) Cumulative food intake measured once per week. C) MRI analysis of lean and fat mass. D) Brown adipose tissue (BAT) weight. E) eWAT weight. F) sWAT weight. G) Hematoxylin and eosin staining of eWAT sections (representative images) and the quantification of adipocyte size. H) Expression of SCD1 and ATGL in liver samples, and FABP4 and leptin in sWAT samples of SF1-*CPT1c*-KO and -WT mice after 8 weeks of HFD. Data are represented as mean ± SEM, male mice, 16-20-week-old, n= 8-10/ group (for H&E staining 2 slices/mouse). *p<0.05 **p<0.01, ***p<0.001, ****p<0.0001. Statistical significance was determined by ANOVA test with post-hoc Bonferroni.

**Fig. 8.**
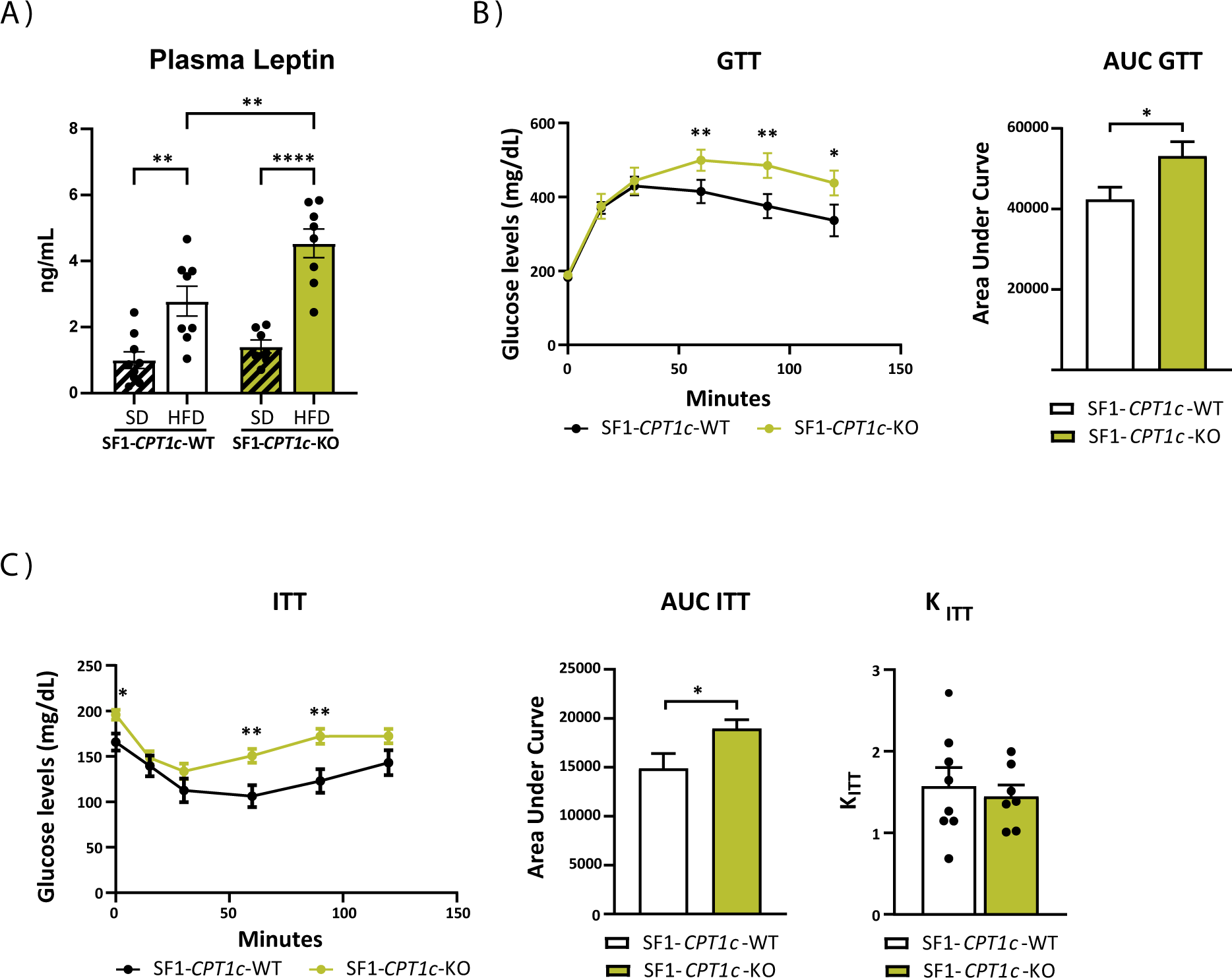
Plasma leptin levels and glucose homeostasis parameters of SF1-*CPT1c*-KO and -WT mice after 8 weeks of HFD. A) Plasma leptin levels. B) Glucose tolerance test (GTT) and quantification of the area under the curve (AUC). C) Insulin tolerance test (ITT) and quantification of the AUC and the K_ITT_ calculated on the first 30 min of ITT. Data were represented as mean ± SEM, male mice, 8-12-week-old, n= 8-10/ group. *p<0.05, **p<0.01, **** p<0.0001. Statistical significance was determined by ANOVA test with post-hoc Bonferroni (GTT and ITT curves) and by t-student test (AUC and K_ITT_).

## DISCUSSION

VMH plays a crucial role in regulating appetite and controlling energy metabolism in the body. Rats with lesions in the VMH suffer transitory hyperphagia and permanent increased adiposity, in addition to insulin resistance (Cox and Powley, 1981). By contrast, an electrical stimulation of the VMH triggers hippophagy, increased EE and a decrease of the RER, suggesting that VMH favors the use of fats as the main energy substrate to supply the cellular oxidation rate in peripheral tissues (Ruffin and Nicolaidis, 1999). Interestingly, intra-VMH injection of a melanocortin receptor agonist (Melanotan II) leads to EE increase and RER decrease, pointing for a role of the melanocortin system in the VMH (Gavini et al., 2016). More recently, low frequency optogenetic or chemogenetic stimulation of SF1 neurons, the most abundant within the VMH, confirmed the contribution of these neurons in regulating feeding, EE and body fat mass independently of locomotor activity and anxiety (Viskaitis et al., 2017; Zhang et al., 2020). Our findings align with these data and identify CPT1c in SF1 neurons as necessary for feeding control and nutrient partitioning in response to a HFD. This is the first report about specific deletion of CPT1c in a hypothalamic nucleus. On the one hand, SF1-*CPT1c*-KO mice, upon exposure to a diet rich in fats, do not appropriately adjust calorie intake and suffer transitory hyperphagia compared to control mice. The impaired expression of orexigenic and anorexigenic neuropeptides and the reduced presence of a-MSH in both the ARC and the PVN are indicative of the inadequate satiety signaling and impaired activation of the melanocortin system. On the other hand, SF1-*CPT1c*-KO mice exhibit a decreased preference for fats as a fuel substrate in the liver and muscle, evidenced by diminished lipolysis and cellular fatty acid oxidation (FAO) in those tissues. Consequently, this leads to increased adiposity and body weight, leptin resistance, and a heightened predisposition to diabetes over the long term. These data also agree with an impaired activation of the melanocortin system, which leads to a decrease in muscle lipolysis and favors WAT expansion in a way independently of food intake (Gavini et al., 2016; Holland et al., 2019; Nogueiras et al., 2007)

It is important to note that these effects are observed in both males and females, albeit slightly less pronounced in females. Notably, CPT1c in SF1 neurons does not appear to play a role in regulating EE or glucose homeostasis, as no differences were observed between genotypes under either SD or short-term HFD conditions. Moreover, no effects on any of the parameters were observed in animals fed a regular chow diet. Hence, CPT1c in SF1 neurons is necessary for adjusting caloric intake and nutrient partitioning in response to an increase in dietary fats, favoring the use of fatty acids as the primary fuel source. However, it is not implicated in the SF1-driven activation of thermogenesis and EE, nor in SF1-mediated recovery from hypoglycemia (Meek et al., 2016). Notably, whole-body CPT1c KO mice did show impaired thermogenesis activation in response to leptin and HFD (Rodríguez-Rodríguez et al., 2019), suggesting the involvement of neurons other than SF1.

This study underscores the important role of CPT1c in sensing dietary fatty acids within the VMH. In control mice, intracerebroventricular injection of oleic acid induces activation of neurons within VMH, while this response is blunted in SF1-*CPT1c*-KO mice. This observation suggests the involvement of CPT1c in the activation of VMH-SF1 neurons and their subsequent signaling to other nuclei that regulate food intake and peripheral fat metabolism. Considering that HFD increases malonyl-CoA levels in the hypothalamus (Tokutake et al., 2012), we hypothesize that SF1 neurons lacking CPT1c fail to sense the changes in malonyl-CoA and consequently do not respond properly to an increase in dietary fats.

What molecular mechanisms might be responsible for these effects? SF1-*CPT1c*-KO mice exhibit notably elevated levels of the two main eCBs, 2-AG and AEA, in hypothalamus under normal conditions, when fed a SD. eCB are retrograde neuromodulators that fine-tune the hypothalamic circuits favoring food intake and energy accumulation (Miralpeix et al., 2021b). Moreover, an elevated tone of the eCB system in the hypothalamus has been associated with an obesogenic phenotype. Consequently, the permanent increase of eCB levels observed in hypothalamus of SF1-*CPT1c*-KO mice could potentially explain the predisposition to weight gain observed in those animals. Additionally, the dynamic of 2-AG in response to the HFD is altered and unlike control mice, there is no observed increase in 2-AG. This difference may be attributed to the already elevated levels of 2-AG in the KO mice. Increased eCBs would also account for the inability of oleic acid to activate VMH neurons, and the disrupted signaling of the melanocortin system upon high fat feeding. We are unaware of the mechanism by which the deficiency in CPT1c increases eCBS levels. Despite we have reported that CPT1c is a regulator of the hydrolase ABHD6 (Miralpeix et al., 2021a), it is well known that the contribution of this enzyme on 2-AG levels in the brain is minimal (Miralpeix et al., 2021b). Hence, further research is necessary to fully comprehend how CPT1c affects the levels of eCBs.

In addition to the eCBs, other CPT1c-down-stream mechanisms might be involved. It is well known that CPT1c regulates neuronal abundance of AMPA receptors, synaptic function and spinogenesis in response to nutrients (Carrasco et al., 2012; Casas et al., 2020; Fadó et al., 2015; Iborra-Lázaro et al., 2023). Therefore, we hypothesize that SF1-neurons of KO mice might not evoke synaptic changes driven by short term fat uptake contributing to metabolic inflexibility.

In summary, results indicate that CPT1c deficiency in SF1 neurons blunts the sensing of dietary fats and increases basal eCBs levels in the hypothalamus. As a consequence, animals do not properly adjust total caloric intake and fuel selection when transitioning from SD to a HFD, leading to impaired fat allocation and obesity development, with the disruption of glucose homeostasis over the long term. These findings shed light on the role of SF1 neurons in fat sensing, regulation of food intake and nutrient partitioning.

## ACKNOWLEDGEMENTS

This study was supported by the Spanish Ministry of Science and Innovation (MCIN) (PID2020-114953RB-C22 to NC, PID2020-114953RB-C21 to LH and DS) co-funded by the European Regional Development Fund, the Biomedical Research Centre in Pathophysiology of Obesity and Nutrition (CIBEROBN) (Grant CB06/03/0001 to LH), the Merck Health Foundation (to LH), the Government of Catalonia (2021SGR00367 to LH) and Xunta de Galicia (predoctoral fellowship to to OF-A; ED481A-2019/026). The authors would also like to acknowledge the INSERM (D.C. and P.Z.), Agence Nationale de la Recherche (ANR-18-CE14-0029, ANR-10-LABX-43 and ANR-10-EQX-008-1 to D.C.), the University of Bordeaux’s IdEx ‘Investments for the Future’ program/GPR BRAIN_2030 (D.C.) and the Fondation pour la Recherche Médicale (EQU202303016291 to D.C.; FRM-SPF202004011774 to C.M).

## AUTOR CONTRIBUTIONS

Conceptualization, A.F., M.L., D.S., D.C., R.R-R. and N.C.; Investigation and Methodology, A.F., D.S.P., S.Z., A.C.R., G.B., M.R-G., J.G-C., C. M., O.F-A., P.Z, L.H.; Software, A.F., D.S.P., S.Z.; Formal Analysis and Data Curation, A.F., D.S.P.; Resources and Funding Acquisition, M.L, D.S., L.H., D.C., R,R-R., and N.C.; Writing – Original Draft Preparation, R.R-R., N.C.; Writing – Review & Editing, all the authors; Supervision, R.R-R. and N.C.

## CONFLICTS OF INTEREST

The authors declare no conflict of interest.

## Notes

### Competing Interest Statement

The authors have declared no competing interest.

## REFERENCES

Cardinal P, André C, Quarta C, Bellocchio L, Clark S, Elie M, Leste-Lasserre T, Maitre M, Gonzales D, Cannich A, Pagotto U, Marsicano G, Cota D. 2014. CB1 cannabinoid receptor in SF1-expressing neurons of the ventromedial hypothalamus determines metabolic responses to diet and leptin. Mol Metab 3. doi:10.1016/j.molmet.2014.07.004

Carrasco P, Sahun I, McDonald J, Ramirez S, Jacas J, Gratacos E, Sierra AY, Serra D, Herrero L, Acker-Palmer A, Hegardt FG, Dierssen M, Casals N. 2012. Ceramide levels regulated by carnitine palmitoyltransferase 1C control dendritic spine maturation and cognition. J Biol Chem 287:21224–21232.

Casals N, Zammit V, Herrero L, Fadó R, Rodríguez-Rodríguez R, Serra D. 2016. Carnitine palmitoyltransferase 1C: From cognition to cancer. Prog Lipid Res 61:134–148. doi:10.1016/j.plipres.2015.11.004

Casas M, Fadó R, Domínguez JL, Roig A, Kaku M, Chohnan S, Solé M, Unzeta M, Miñano-Molina AJ, Rodríguez-Álvarez J, Dickson EJ, Casals N. 2020. Sensing of nutrients by CPT1C controls SAC1 activity to regulate AMPA receptor trafficking. J Cell Biol 219. doi:10.1083/jcb.201912045

Castellanos-Jankiewicz A, Guzmán-Quevedo O, Fénelon VS, Zizzari P, Quarta C, Bellocchio L, Tailleux A, Charton J, Fernandois D, Henricsson M, Piveteau C, Simon V, Allard C, Quemener S, Guinot V, Hennuyer N, Perino A, Duveau A, Maitre M, Leste-Lasserre T, Clark S, Dupuy N, Cannich A, Gonzales D, Deprez B, Mithieux G, Dombrowicz D, Bäckhed F, Prevot V, Marsicano G, Staels B, Schoonjans K, Cota D. 2021. Hypothalamic bile acid-TGR5 signaling protects from obesity. Cell Metab 33:1483–1492.e10. doi:10.1016/J.CMET.2021.04.009

Cha SH, Rodgers JT, Puigserver P, Chohnan S, Lane MD. 2006. Hypothalamic malonyl-CoA triggers mitochondrial biogenesis and oxidative gene expression in skeletal muscle: Role of PGC-1alpha. Proc Natl Acad Sci U S A 103:15410–15415.

Cox JE, Powley TL. 1981. Prior vagotomy blocks VMH obesity in pair-fed rats. Am J Physiol Endocrinol Metab 3. doi:10.1152/ajpendo.1981.240.5.e573

Dai Y, Wolfgang MJ, Cha SH, Lane MD. 2007. Localization and effect of ectopic expression of CPT1c in CNS feeding centers. Biochem Biophys Res Commun 359:469–474.

Fadó R, Rodríguez-Rodríguez R, Casals N. 2021. The return of malonyl-CoA to the brain: Cognition and other stories. Prog Lipid Res. doi:10.1016/j.plipres.2020.101071

Fadó R, Soto D, Miñano-Molina AJ, Pozo M, Carrasco P, Yefimenko N, Rodríguez-Álvarez J, Casals N. 2015. Novel Regulation of the Synthesis of α-Amino-3-hydroxy-5-methyl-4-isoxazolepropionic Acid (AMPA) Receptor Subunit GluA1 by Carnitine Palmitoyltransferase 1C (CPT1C) in the Hippocampus. J Biol Chem 290:25548–60. doi:10.1074/jbc.M115.681064

Fosch A, Rodriguez-Garcia M, Miralpeix C, Zagmutt S, Larrañaga M, Reguera AC, Garcia-Chica J, Herrero L, Serra D, Casals N, Rodriguez-Rodriguez R. 2023. Central Regulation of Brown Fat Thermogenesis in Response to Saturated or Unsaturated Long-Chain Fatty Acids. Int J Mol Sci 24. doi:10.3390/ijms24021697

Fosch A, Zagmutt S, Casals N, Rodríguez-Rodríguez R. 2021. New insights of sf1 neurons in hypothalamic regulation of obesity and diabetes. Int J Mol Sci 22. doi:10.3390/ijms22126186

Gao L, Chiou W, Tang H, Cheng X, Camp HS, Burns DJ. 2007. Simultaneous quantification of malonyl-CoA and several other short-chain acyl-CoAs in animal tissues by ion-pairing reversed-phase HPLC/MS. Journal of Chromatography B 853:303–313. doi:10.1016/j.jchromb.2007.03.029

Gao S, Zhu G, Gao X, Wu D, Carrasco P, Casals N, Hegardt FG, Moran TH, Lopaschuk GD. 2011. Important roles of brain-specific carnitine palmitoyltransferase and ceramide metabolism in leptin hypothalamic control of feeding. Proc Natl Acad Sci U S A 108:9691–9696.

Gavini CK, Jones WC, Novak CM. 2016. Ventromedial hypothalamic melanocortin receptor activation: regulation of activity energy expenditure and skeletal muscle thermogenesis. Journal of Physiology 594. doi:10.1113/JP272352

Gratacòs-Batlle E, Olivella M, Sánchez-Fernández N, Yefimenko N, Miguez-Cabello F, Fadó R, Casals N, Gasull X, Ambrosio S, Soto D. 2018. Mechanisms of CPT1C-Dependent AMPAR Trafficking Enhancement. Front Mol Neurosci 11:275. doi:10.3389/fnmol.2018.00275

He W, Lam TK, Obici S, Rossetti L. 2006. Molecular disruption of hypothalamic nutrient sensing induces obesity. Nat Neurosci 9:227–233.

Holland J, Sorrell J, Yates E, Smith K, Arbabi S, Arnold M, Rivir M, Morano R, Chen J, Zhang X, Dimarchi R, Woods SC, Sanchez-Gurmaches J, Wohleb E, Perez-Tilve D. 2019. A Brain-Melanocortin-Vagus Axis Mediates Adipose Tissue Expansion Independently of Energy Intake. Cell Rep 27. doi:10.1016/j.celrep.2019.04.089

Hu Z, Cha SH, Chohnan S, Lane MD. 2003. Hypothalamic malonyl-CoA as a mediator of feeding behavior. Proceedings of the National Academy of Sciences 100:12624– 12629. doi:10.1073/pnas.1834402100

Iborra-Lázaro G, Djebari S, Sánchez-Rodríguez I, Gratacòs-Batlle E, Sánchez-Fernández N, Radošević M, Casals N, Navarro-López J de D, Soto del Cerro D, Jiménez-Díaz L. 2023. CPT1C is required for synaptic plasticity and oscillatory activity that supports motor, associative and non-associative learning. Journal of Physiology 601. doi:10.1113/JP284248

Ikeda Y, Luo X, Abbud R, Nilson JH, Parker KL. 1995. The nuclear receptor steroidogenic factor 1 is essential for the formation of the ventromedial hypothalamic nucleus. Molecular Endocrinology 9. doi:10.1210/mend.9.4.7659091

Kim KW, Jo YH, Zhao L, Stallings NR, Chua SC, Parker KL. 2008. Steroidogenic factor 1 regulates expression of the cannabinoid receptor 1 in the ventromedial hypothalamic nucleus. Molecular Endocrinology 22. doi:10.1210/me.2008-0127

Kim KW, Zhao L, Donato J, Kohno D, Xu Y, Eliasa CF, Lee C, Parker KL, Elmquist JK. 2011. Steroidogenic factor 1 directs programs regulating diet-induced thermogenesis and leptin action in the ventral medial hypothalamic nucleus. Proc Natl Acad Sci U S A 108. doi:10.1073/pnas.1102364108

Kim KW, Zhao L, Parker KL. 2009. Central nervous system-specific knockout of steroidogenic factor 1. Mol Cell Endocrinol. doi:10.1016/j.mce.2008.09.026

Lane MD, Wolfgang M, Cha S-H, Dai Y. 2008. Regulation of food intake and energy expenditure by hypothalamic malonyl-CoA. Int J Obes (Lond) 32 **Suppl 4**:S49–54. doi:10.1038/ijo.2008.123

Liu Y, Colby JK, Zuo X, Jaoude J, Wei D, Shureiqi I. 2018. The Role of PPAR-δ in Metabolism, Inflammation, and Cancer: Many Characters of a Critical Transcription Factor. Int J Mol Sci 19. doi:10.3390/IJMS19113339

López M, Lelliott CJ, Tovar S, Kimber W, Gallego R, Virtue S, Blount M, Vázquez MJ, Finer N, Powles TJ, O’Rahilly S, Saha AK, Diéguez C, Vidal-Puig AJ. 2006. Tamoxifen-induced anorexia is associated with fatty acid synthase inhibition in the ventromedial nucleus of the hypothalamus and accumulation of malonyl-CoA. Diabetes 55:1327–1336. doi:10.2337/db05-1356

Meek TH, Nelson JT, Matsen ME, Dorfman MD, Guyenet SJ, Damian V, Allison MB, Scarlett JM, Nguyen HT, Thaler JP, Olson DP, Myers MG, Schwartz MW, Morton GJ. 2016. Functional identification of a neurocircuit regulating blood glucose. Proc Natl Acad Sci U S A 113. doi:10.1073/pnas.1521160113

Miralpeix C, Fosch A, Casas J, Baena M, Herrero L, Serra D, Rodríguez-Rodríguez R, Casals N. 2019. Hypothalamic endocannabinoids inversely correlate with the development of diet-induced obesity in male and female mice. J Lipid Res 60. doi:10.1194/jlr.M092742

Miralpeix C, Reguera AC, Fosch A, Casas M, Lillo J, Navarro G, Franco R, Casas J, Alexander SPH, Casals N, Rodríguez-Rodríguez R. 2021a. Carnitine palmitoyltransferase 1C negatively regulates the endocannabinoid hydrolase ABHD6 in mice, depending on nutritional status. Br J Pharmacol 178. doi:10.1111/bph.15377

Miralpeix C, Reguera AC, Fosch A, Zagmutt S, Casals N, Cota D, Rodríguez-Rodríguez R. 2021b. Hypothalamic endocannabinoids in obesity: an old story with new challenges. Cellular and Molecular Life Sciences. doi:10.1007/s00018-021-04002-6

Nogueiras R, Wiedmer P, Perez-Tilve D, Veyrat-Durebex C, Keogh JM, Sutton GM, Pfluger PT, Castaneda TR, Neschen S, Hofmann SM, Howles PN, Morgan DA, Benoit SC, Szanto I, Schrott B, Schürmann A, Joost HG, Hammond C, Hui DY, Woods SC, Rahmouni K, Butler AA, Farooqi IS, O’Rahilly S, Rohner-Jeanrenaud F, Tschöp MH. 2007. The central melanocortin system directly controls peripheral lipid metabolism. Journal of Clinical Investigation 117. doi:10.1172/JCI31743

Okamoto S, Sato T, Tateyama M, Kageyama H, Maejima Y, Nakata M, Hirako S, Matsuo T, Kyaw S, Shiuchi T, Toda C, Sedbazar U, Saito K, Asgar NF, Zhang B, Yokota S, Kobayashi K, Foufelle F, Ferré P, Nakazato M, Masuzaki H, Shioda S, Yada T, Kahn BB, Minokoshi Y. 2018. Activation of AMPK-Regulated CRH Neurons in the PVH is Sufficient and Necessary to Induce Dietary Preference for Carbohydrate over Fat. Cell Rep 22:706–721. doi:10.1016/j.celrep.2017.11.102

Palomo-Guerrero M, Fadó R, Casas M, Pérez-Montero M, Baena M, Helmer PO, Domínguez JL, Roig A, Serra D, Hayen H, Stenmark H, Raiborg C, Casals N. 2019. Sensing of nutrients by CPT1C regulates late endosome/lysosome anterograde transport and axon growth. Elife 8. doi:10.7554/eLife.51063

Pozo M, Rodríguez-Rodríguez R, Ramírez S, Seoane-Collazo P, López M, Serra D, Herrero L, Casals N. 2017. Hypothalamic Regulation of Liver and Muscle Nutrient Partitioning by Brain-Specific Carnitine Palmitoyltransferase 1C in Male Mice. Endocrinology 158:2226–2238. doi:10.1210/en.2017-00151

Price N, van der Leij F, Jackson V, Corstorphine C, Thomson R, Sorensen A, Zammit V. 2002. A novel brain-expressed protein related to carnitine palmitoyltransferase I. Genomics 80:433–442.

Price Nigel, van der Leij F, Jackson V, Corstorphine C, Thomson R, Sorensen A, Zammit V. 2002. A novel brain-expressed protein related to carnitine palmitoyltransferase I. Genomics 80:433–442. doi:10.1006/geno.2002.6845

Ramírez S, Martins L, Jacas J, Carrasco P, Pozo M, Clotet J, Serra D, Hegardt FG, Diéguez C, López M, Casals N. 2013. Hypothalamic ceramide levels regulated by CPT1C mediate the orexigenic effect of ghrelin. Diabetes 62:2329–37. doi:10.2337/db12-1451

Rodríguez-Rodríguez R, Fosch A, Garcia-Chica J, Zagmutt S, Casals N. 2023. Targeting carnitine palmitoyltransferase 1 isoforms in the hypothalamus: A promising strategy to regulate energy balance. J Neuroendocrinol 35. doi:10.1111/jne.13234

Rodríguez-Rodríguez R, Miralpeix C, Fosch A, Pozo M, Calderón-Domínguez M, Perpinyà X, Vellvehí M, López M, Herrero L, Serra D, Casals N. 2019. CPT1C in the ventromedial nucleus of the hypothalamus is necessary for brown fat thermogenesis activation in obesity. Mol Metab 19:75–85. doi:10.1016/j.molmet.2018.10.010

Ruffin MP, Nicolaidis S. 1999. Electrical stimulation of the ventromedial hypothalamus enhances both fat utilization and metabolic rate that precede and parallel the inhibition of feeding behavior. Brain Res 846. doi:10.1016/S0006-8993(99)01922-8

Tokutake Y, Iio W, Onizawa N, Ogata Y, Kohari D, Toyoda A, Chohnan S. 2012. Effect of diet composition on coenzyme A and its thioester pools in various rat tissues. Biochem Biophys Res Commun 423:781–784. doi:10.1016/j.bbrc.2012.06.037

Tokutake Y, Onizawa N, Katoh H, Toyoda A, Chohnan S. 2010. Coenzyme A and its thioester pools in fasted and fed rat tissues. Biochem Biophys Res Commun 402:158–162. doi:10.1016/j.bbrc.2010.10.009

Viskaitis P, Irvine EE, Smith MA, Choudhury AI, Alvarez-Curto E, Glegola JA, Hardy DG, Pedroni SMA, Paiva Pessoa MR, Fernando ABP, Katsouri L, Sardini A, Ungless MA, Milligan G, Withers DJ. 2017. Modulation of SF1 Neuron Activity Coordinately Regulates Both Feeding Behavior and Associated Emotional States. Cell Rep 21. doi:10.1016/j.celrep.2017.11.089

Wang H, Eckel RH. 2009. Lipoprotein lipase: from gene to obesity. Am J Physiol Endocrinol Metab 297. doi:10.1152/AJPENDO.90920.2008

Wolfgang MJ, Cha SH, Sidhaye A, Chohnan S, Cline G, Shulman GI, Lane MD. 2007. Regulation of hypothalamic malonyl-CoA by central glucose and leptin. Proc Natl Acad Sci U S A 104:19285–90. doi:10.1073/pnas.0709778104

Wolfgang MJ, Kurama T, Dai Y, Suwa A, Asaumi M, Matsumoto S, Cha SH, Shimokawa T, Lane MD. 2006. The brain-specific carnitine palmitoyltransferase-1c regulates energy homeostasis. Proc Natl Acad Sci U S A 103:7282–7287.

Zagmutt S, Mera P, González-García I, Ibeas K, Romero M del M, Obri A, Martin B, Esteve-Codina A, Soler-Vázquez MC, Bastias-Pérez M, Cañes L, Augé E, Pelegri C, Vilaplana J, Ariza X, García J, Martinez-González J, Casals N, López M, Palmiter R, Sanz E, Quintana A, Herrero L, Serra D. 2023. CPT1A in AgRP neurons is required for sex-dependent regulation of feeding and thirst. Biol Sex Differ 14. doi:10.1186/s13293-023-00498-8

Zhang J, Chen D, Sweeney P, Yang Y. 2020. An excitatory ventromedial hypothalamus to paraventricular thalamus circuit that suppresses food intake. Nat Commun 11. doi:10.1038/s41467-020-20093-4

