## Supplementary data for "DEFICIENCY OF THE NUTRIENT SENSOR CPT1c IN SF1 NEURONS DISRUPTS THE ENDOCANNABINOID SYSTEM RESULTING IN COMPROMISED SATIETY AND FUEL SELECTION UPON FAT INTAKE"

### Supplemental figures

Fosch et al, 2023

A)

| GENOTYPE | PRIMER | SEQUENCE | AMPLICON SIZE | VOLUME | AGAROSE GEL | ANNEALING |
|---|---|---|---|---|---|---|
| CPT1c – KO | F9 | 5'- GAGTCAGCCATGACCCGACTGTT -3' | 200pb | 0.9 µL | 0.8% | 65ºC – 30'' |
| | R1 | 5'- CCGGTAGAATTGACCTGCAGGGGC -3' | | 0.6 µL | | |
| | R9 | 5'- CGCTAAAGCCCAGACAGAACACAC -3' | | 1.6 µL | | |
| KO FIRST 1 | F – 5'arm | 5'- ATGGACCCATAGTCTTGAAACCTGG-3' | 625 pb and 432pb | 2 µL | 2% | 63.5ºC – 30'' |
| | F - Neo | 5'- GGGATCTCATGCTGGAGTTCTTCG -3' | | | | |
| | R - Targ | 5'- GTGTTTGCCTAGCATGCGAACG -3' | | | | |
| KO FIRST 2 | F - Lox | 5'- GAGATGGCGCAACGCAATTAATG -3' | 292bp | 2 µL | 2% | 63.5ºC – 30'' |
| | R – 3'arm | 5'- ATTCTTGATTCGTGTGACTCATTCCG -3' | | | | |
| CONDITIONAL KO/FLOXED | F – 5'arm | 5'- ATGGACCCATAGTCTTGAAACCTGG-3' | 517pb | 1.5 µL | 2% | 63.5ºC – 30'' |
| | R - Targ | 5'- GTGTTTGCCTAGCATGCGAACG -3' | | | | |
| FLPO | F – FlpO | 5'- CTATCGAATTCCACCATGGCTCCTAAGAAGAA -3' | 1300pb | 1 µL | 0.8% | 58ºC – 1' |
| | R - FlpO | 5'- CAATGCGATGAATTCTCAGATCCGCCTCTTGATGTA -3' | | | | |
| SF1-CRE | F – Cre | 5'- CTGAGCTGCAGCGCAGGGACAT -3' | 250pb | 1 µL | 2% | 68ºC – 1' |
| | R – Cre | 5'-TGCGAACCTCATCACTCGTTGCAT -3' | | | | |

B)

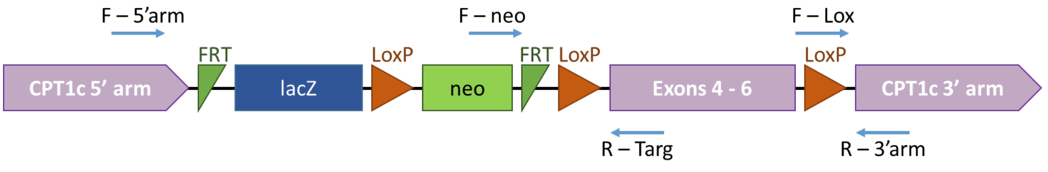


**Supplemental Figure 1. Genotyping primers.** A) Oligonucleotides enhancers used for genotyping mice. Sequence and experimental details are also indicated. B) Schematic representation of modified Cpt1c gene, and all oligonucleotide enhancers used for genotyping.

| Gene (protein) | Forward sequence | Reverse sequence |
|---|---|---|
| *Acaca (ACC$\alpha$)* | 5'-ATGGGCGGAATGGTCTCTTTC | 5'-TGGGGACCTTGTCTTCATCAT |
| *Agrp* | 5'-TTTGTCCTCTGAAGCTGTATGC | 5'-GCATGAGGTGCCTCCCTA |
| *Atgl* | 5'-AACAACAGCATCCAGTTCAA | 5'-GGTTCAGTAGGTCATTCCTC |
| *Cd36* | 5'-TGGCCAAGCTATTGCGACAT | 5'-ACACAGCGTAGATAGACCTGC |
| *Cpt1a* | 5'-GACTCCGCTCGCTCATTC | 5'-AAGGCCACAGCTTGGTGA |
| *Cpt1b* | 5'-TGCCTTTACATCGTCTCCAA | 5'-GGCTCCAGGGTTCAGAAAGT |
| *Dagla* | 5'-TATCTTCCTCTTCCTGCT | 5'-CCATTTCGGCAATCATAC |
| *Faah* | 5'-CAGCTACAAGGGCCATGCT | 5'-TTCCACGGGTTCATGGTCTG |
| *Fabp4* | 5'-GGATGGAAAGTCGACCACAA | 5'-TGGAAGTCACGCCTTTCATA |
| *Fasn* | 5'-CAGATGATGACAGGAGATGGAA | 5'-CACTCACACCCACCCAGA |
| *Gapdh* | 5'-TCCACTTTGCCACTGCA | 5'-GAGACGGCCGCATCTTCTT |
| *Hsl* | 5'-TCACGCTACATAAAGGCTGCT | 5'-CCACCCGTAAAGAGGGAACT |
| *Leptin* | 5'-CAGGATCAATGACATTTCACACA | 5'-GCTGGTGAGGACCTGTTGAT |
| *Lpl* | 5'-GGGAGTTTGGCTCCAGAGTTT | 5'-TGTGTCTTCAGGGGTCCTTAG |
| *Mgl1* | 5'-CCCAGTGGCACACCCAAG | 5'-TAACGGCCACAGTGTTCCC |
| *Nape-pld* | 5'-AAAACATCTCCATCCCGAA | 5'-CGTCCATTTCCACCATCA |
| *Npy* | 5'-TCCGCTCTGCGACACTACAT | 5'-TGCTTTCCTTCATTAAGAGGT |
| *Pgc1a* | 5'-GAAAGGGCCAAACAGAGAGA | 5'-GTAAATCACACGGCGCTCTT |
| *Pomc* | 5'-TGAACATCTTTGTCCCCAGAG | 5'-TGCAGAGGCAAACAAGATTGG |
| *Ppard* | 5'-AGGAGAAAGAGGAAGTGGCC | 5'-GGGAGGAATTCTGGGAGAGG |
| *Pparg* | 5'-CCGAAGAACCATCCGATTGAA | 5'-GCCCAAACCTGATGGCATT |
| *Prdm6* | 5'-CCTAAGGTGTGCCCAGCA | 5'-CACCTTCCGCTTTTCTACCC |
| *Scd1* | 5'-TTCCCTCCTGCAAGCTCTAC | 5'-CAGAGCGCTGGTCATGTAGT |
| *Adrb3 ($\beta$3AR)* | 5'-TCGACATGTTCCTCCACAAA | 5'-GATGGTCCAAGATGGTGCTT |

**Supplemental Figure 2. Oligonucleotides used for quantitative RT-PCR**

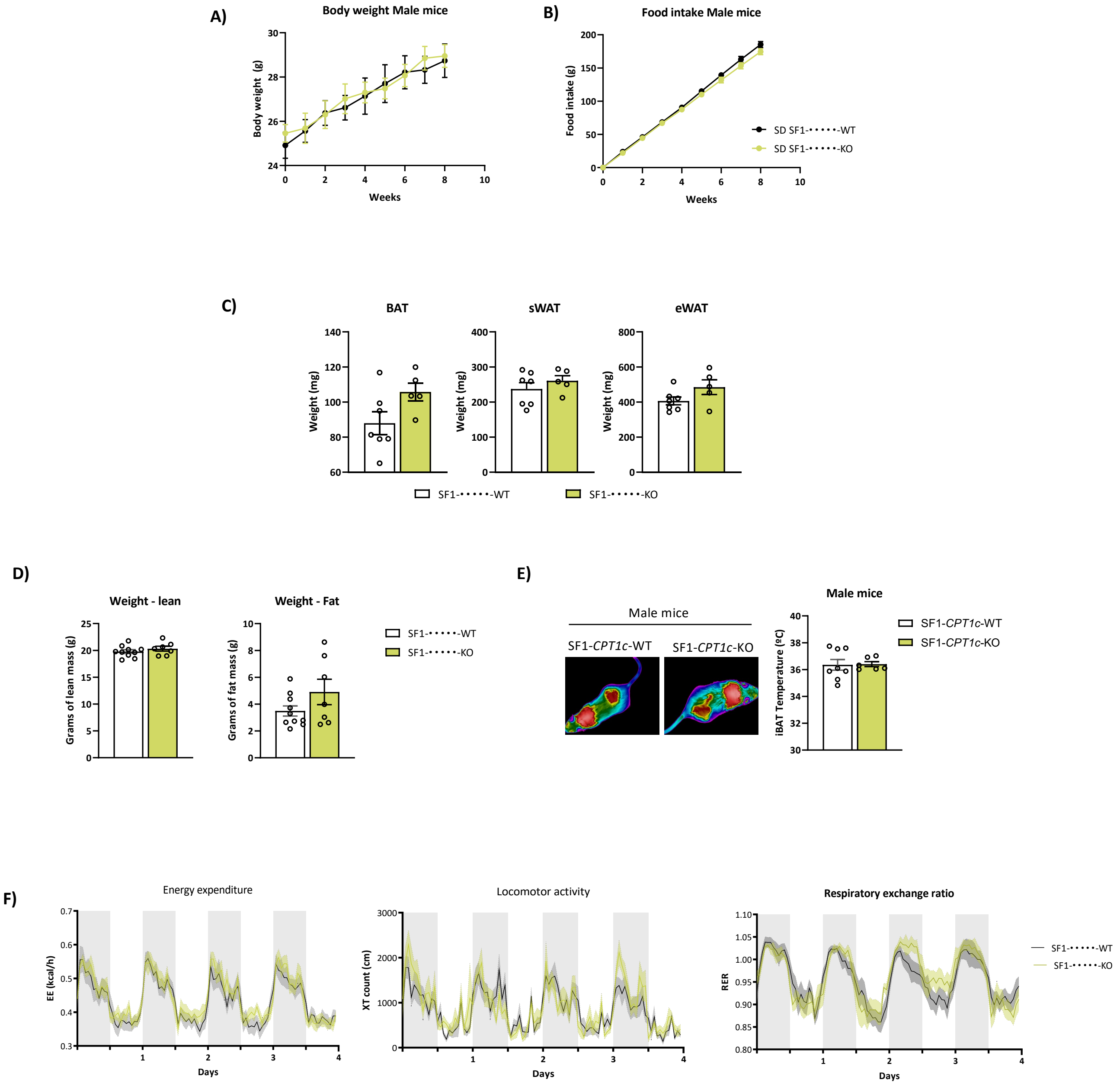


**Supplemental Figure 3. Metabolic phenotype of SF1-CPT1c-KO male mice under chow diet. A)** Body weight. B) Food intake. C) Weight of BAT, sWAT and eWAT collected after 8 weeks on chow diet. D) MRI analysis of lean and fat mass. E) Representative IR pictures and iBAT temperature normalized to the temperature of back's mice. F) Energy expenditure, locomotor activity on the horizontal axis of the cage (XT) and Respiratory exchange ratio monitored for 4 days. Data were represented as mean ± SEM, male mice, between 8 and 12-week-old, n= 7-10/ group. Statistical significance was determined by t-student test and by ANOVA test with post-hoc Bonferroni.

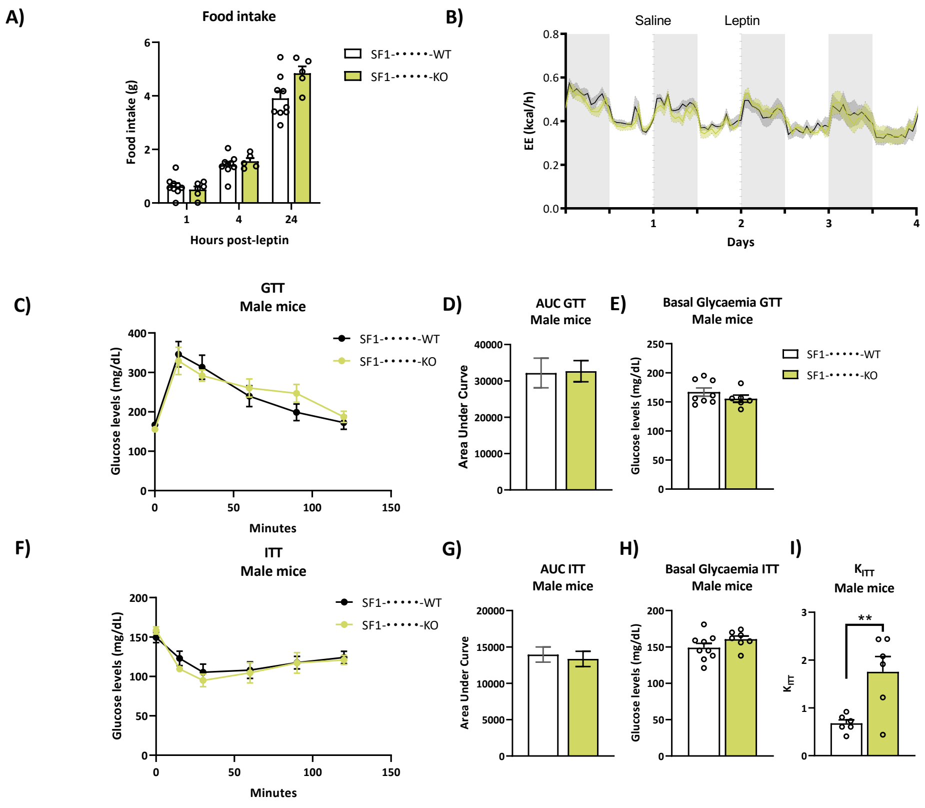


**Supplemental Figure 4. Leptin and insulin sensitivity of male mice in chow diet**. A) Leptin sensitivity test. Food intake post-leptin ip injection. B) Energy expenditure (EE) after leptin ip injection. Data were represented as mean ± SEM, 15-week-old male mice, n= 7-10/ group. Statistical significance was determined by ANOVA test with post-hoc Bonferroni. C) GTT. D) Area under curve of GTT. E) Basal glycaemia of GTT. F) ITT. G) Area under curve of ITT. H) Basal glycaemia of ITT. I) KITT calculated on the first 30 min of ITT. Data were represented as mean ± SEM, male mice, 8-weekold, n= 7-9/ group. Statistical significance was determined by ANOVA test and t-student test.

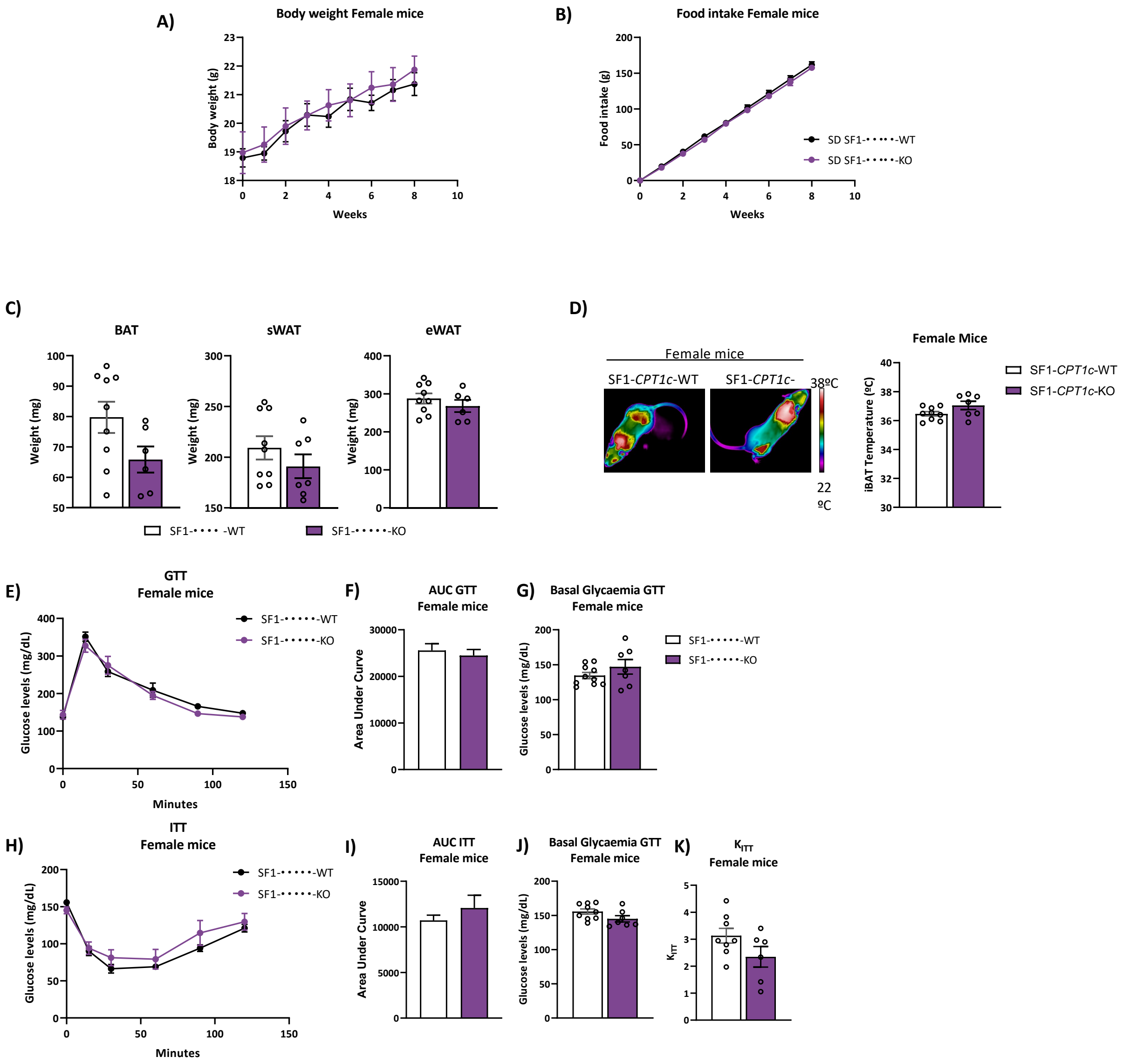


**Supplemental Figure 5. Metabolic phenotype of SF1-CPT1c-KO female mice under chow diet. A)** Body weight. B) Food intake. C) Weight of BAT, sWAT and eWAT collected after 8 weeks on chow diet. D) Representative IR pictures and iBAT temperature normalized to the temperature of back's mice. E) GTT. F) Area under curve of GTT. G) Basal glycaemia of GTT.H) ITT. I) Area under curve of ITT. J) Basal glycaemia of ITT. K) KITT calculated on the first 30 min of ITT. Data were represented as mean ± SEM, male mice, 8-weeks-old, n= 7-9/ group. Statistical significance was determined by ANOVA test and t-student test.

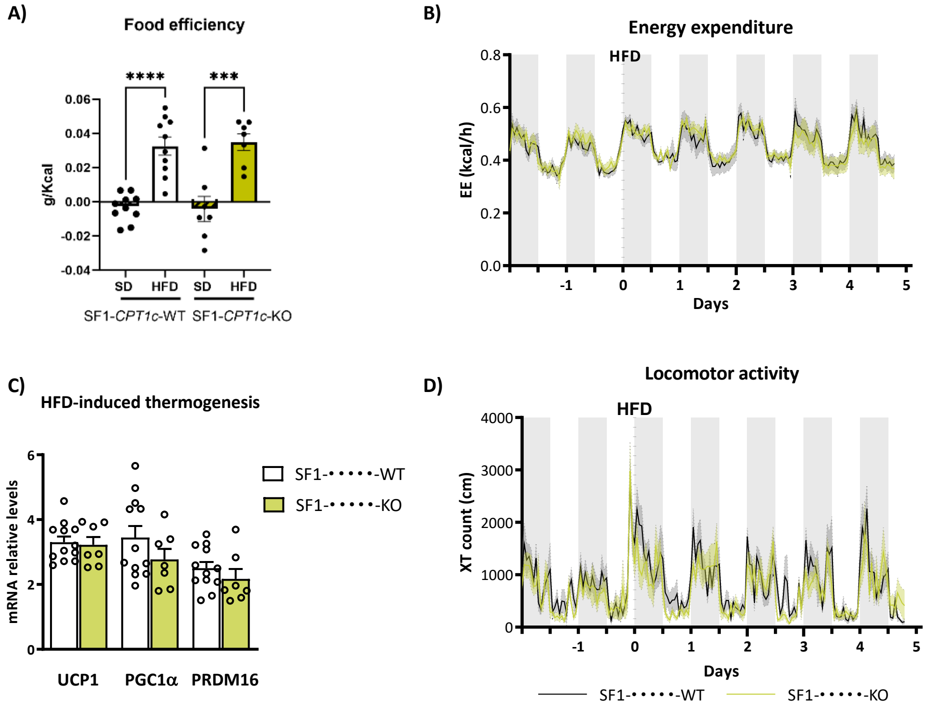


**Supplemental Figure 6. Short-term high fat diet (HFD) in male mice.** A) Food efficiency, calculated by dividing the weight gained between the kilocalories consumed. n=7-10, One Way ANOVA. B) energy expenditure. C) UCO1, PGC1α and PRDM16 mRNA thermogenic markers. D) Locomotor activity on the horizontal axis of the cage (XT). Data were represented as mean ± SEM, male mice, between 16-week-old, n= 7-10/ group. Statistical significance was determined by ANOVA test with post-hoc Bonferroni.

A)

B)

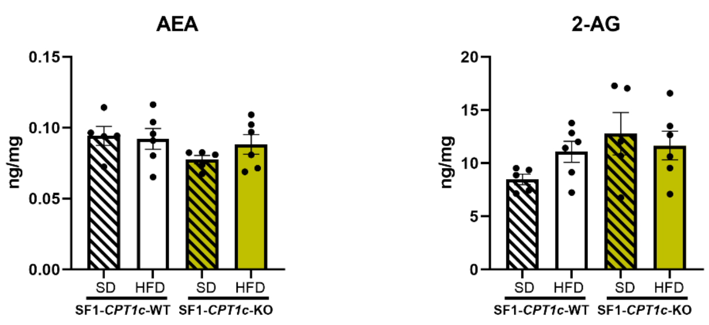


**Supplemental Figure 7: eCB levels in the hippocampus of SF1-Cpt1c-KO and -WT mice under standard diet (SD) or HFD.** A) Anandamide (AEA), B) 2-arachidonoylglycerol (2-AG). Data are represented as mean ± SEM, male mice, 8-12 week-old, n= 5-6/group. No statistical significance is observed. Statistical significance was determined by ANOVA test.

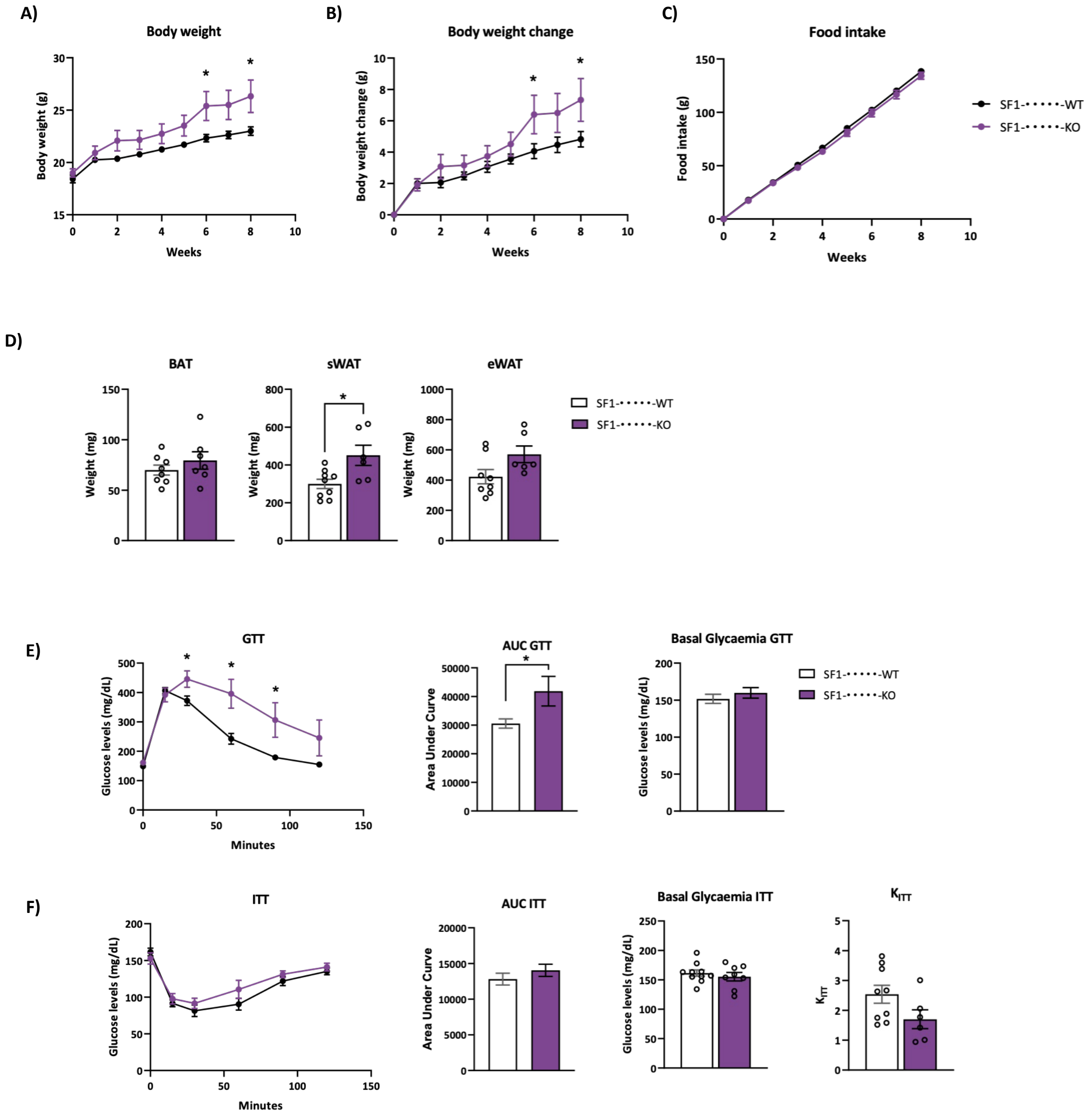


**Supplemental Figure 8. Metabolic phenotype of SF1-CPT1c-KO female mice under 8 weeks of HFD exposure.** A) Body weight. B) Body weight change. C) Food intake. D) Weight of BAT, sWAT and eWAT collected after 8 weeks on high-fat diet. E) GTT. F) Area under curve of GTT. G) Basal glycaemia of GTT.H) ITT. I) Area under curve of ITT. J) Basal glycaemia of ITT. K) KITT calculated on the first 30 min of ITT. All tests performed after 8 weeks of high-fat diet exposure. Data were represented as mean ± SEM, male mice, 8-weeks-old, n= 7-9/ group. Statistical significance was determined by ANOVA test and t-student test.
